# Lipid Sponge Droplets as Programmable Synthetic Organelles

**DOI:** 10.1101/2020.01.20.913350

**Authors:** Ahanjit Bhattacharya, Henrike Niederholtmeyer, Kira A. Podolsky, Rupak Bhattacharya, Jing-Jin Song, Roberto J. Brea, Chu-Hsien Tsai, Sunil K. Sinha, Neal K. Devaraj

## Abstract

Living cells segregate molecules and reactions in various subcellular compartments known as organelles. Spatial organization is likely essential for expanding the biochemical functions of synthetic reaction systems, including artificial cells. Many studies have attempted to mimic organelle functions using lamellar membrane-bound vesicles. However, vesicles typically suffer from highly limited transport across the membranes and an inability to mimic the dense membrane networks typically found in organelles such as the endoplasmic reticulum. Here we describe programmable synthetic organelles based on highly stable nonlamellar sponge phase droplets that spontaneously assemble from a single-chain galactolipid and non-ionic detergents. Due to their nanoporous structure, lipid sponge droplets readily exchange materials with the surrounding environment. In addition, the sponge phase contains a dense network of lipid bilayers and nanometric aqueous channels, which allows different classes of molecules to partition based on their size, polarity, and specific binding motifs. The sequestration of biologically relevant macromolecules can be programmed by the addition of suitably functionalized amphiphiles to the droplets. We demonstrate that droplets can harbor functional soluble and transmembrane proteins, allowing for the co-localization and concentration of enzymes and substrates to enhance reaction rates. Droplets protect bound proteins from proteases, and these interactions can be engineered to be reversible and optically controlled. Our results show that lipid sponge droplets permit the facile integration of membrane-rich environments and self-assembling spatial organization with biochemical reaction systems.

**Significance statement:** Organelles spatially and temporally orchestrate biochemical reactions in a cell to a degree of precision that is still unattainable in synthetic reaction systems. Additionally, organelles such as the endoplasmic reticulum (ER) contain highly interconnected and dense membrane networks that provide large reaction spaces for both transmembrane and soluble enzymes. We present lipid sponge droplets to emulate the functions of organelles such as the ER. We demonstrate that lipid sponge droplets can be programmed to internally concentrate specific proteins, host and accelerate biochemical transformations, and to rapidly and reversibly sequester and release proteins to control enzymatic reactions. The self-assembled and programmable nature of lipid sponge droplets will facilitate the integration of complex functions for bottom up synthetic biology.

## Introduction

Compartmentalization is a defining feature of life. Eukaryotic cells are characterized by their membrane-bound and membraneless organelles (1) and in prokaryotic cells, different processes are precisely organized in subcellular locations and microcompartments (2). Spatial organization allows cells to co-localize and concentrate molecules, establish concentration gradients, separate incompatible reactions, and regulate reactions by controlling the availability of reactants. The biochemical capabilities of “bottom up” cell-mimics and synthetic reaction systems lag behind those of living cells, in part due to a lack of programmable compartmentalization capabilities.

Several methodologies have been developed to sequester specific biochemical processes to defined locations and to emulate functions of natural cellular organelles (3). Examples include the use of coacervates (4–6), hydrogels (7–9), as well as protein (10, 11) and polymer (12) assemblies to spatially organize biomolecules. In eukaryotic cells, compartmentalization is primarily achieved by lipid bilayer membranes that demarcate the boundary to the outside world and enclose classical organelles. Lipid bilayers serve as the necessary matrix for the proper folding and function of transmembrane proteins, can scaffold protein signaling complexes, and provide an interface for sequestering molecules and proteins from the environment. As such, lamellar lipid membrane-bound vesicles have been extensively utilized as experimental model systems to emulate the plasma membrane (13) as well as that of organelles like chloroplasts (14, 15) and nuclei (16).

While lamellar vesicles are conceptually attractive as membrane mimics, transport of molecules is severely restricted due to the impermeable nature of lipid bilayers. Cells work around this limitation through the incorporation of highly sophisticated transport machinery, including membrane channels, molecular transporters, and intracellular vesicular trafficking. The reconstitution of such systems in vitro for the design of artificial organelles is highly challenging. In addition, membrane-bound vesicles typically fail to capture the high density of lipid bilayer networks present in many natural cellular organelles such as the endoplasmic reticulum (ER), Golgi apparatus, mitochondria and chloroplasts. These dense membrane networks offer enormous surface area for membrane-associated reactions. In eukaryotic cells, the ER is often the organelle with the greatest surface area and may contain 50-60% of the total membrane (17). Lipid membranes of the ER form a vast convoluted network interspersed by aqueous channels. Combined, the issues of restricted transport and limited membrane surface area have hindered efforts to use lamellar vesicles to create effective lipid-containing structures that act as bioreactors mimicking the function of organelles.

In contrast to the closed lamellar lipid bilayers found in membrane-bound vesicles, bicontinuous lipidic materials such as sponge phases (also referred to as L3 phase or lipid coacervates) are inherently open structures and possess properties that resemble highly membranous organelles such as the ER. They contain a highly convoluted and interconnected three dimensional network of lipid bilayers intersected by similarly interconnected nanometer sized aqueous channels (18, 19). Bicontinuous lipid phases (sponge and cubic) offer enormous membrane surface areas, the capacity to harbor both hydrophobic and hydrophilic molecules, and the ability to sequester soluble and membrane proteins (20).

We hypothesized that compartments consisting of bicontinuous lipid sponge phases could serve as ideal membrane matrices for organizing biochemical reactions. This is due to their ability to both readily take up and release molecules from the environment and their high surface area. However, while sponge phases have been observed in a variety of surfactant systems, previously described systems frequently form a single bulk phase, exist only within a narrow range of concentration, pH, and temperature, and are highly sensitive to additives (19, 21–24). Here we show that a combination of single-chain galactolipids such as that *N*-oleoyl β-D-galactopyranosylamine (GOA) and commercially available non-ionic surfactants such as octylphenoxypolyethoxyethanol (IGEPAL, **Fig. 1*A***) form micron-sized spherical droplets in aqueous media that consist of a highly stable and biocompatible lipid sponge phase. Using a combination of physical and biochemical techniques, including X-ray diffraction and cryo-electron microscopy, we have characterized the nonlamellar structure of the lipid sponge droplets. We show that uptake of specific biomolecules into the droplets can be predictably programmed. Importantly, the dense lipid bilayer network within the droplets allows spontaneous reconstitution of functional transmembrane proteins, emulating the biosynthetic functions of the endomembrane system. To mimic additional organelle functions, we demonstrate that co-localization and concentration of an enzyme and its substrate in droplets enhances the rate of a biochemical reactions, and that reversible light-controlled sequestration and release of a protein can be used to regulate enzymatic reactivity. By providing a nonlamellar alternative to membrane-bound vesicles, lipid sponge droplets offer a unique platform for mimicking both membrane- and solution-associated organelle functions.

**Fig. 1.**
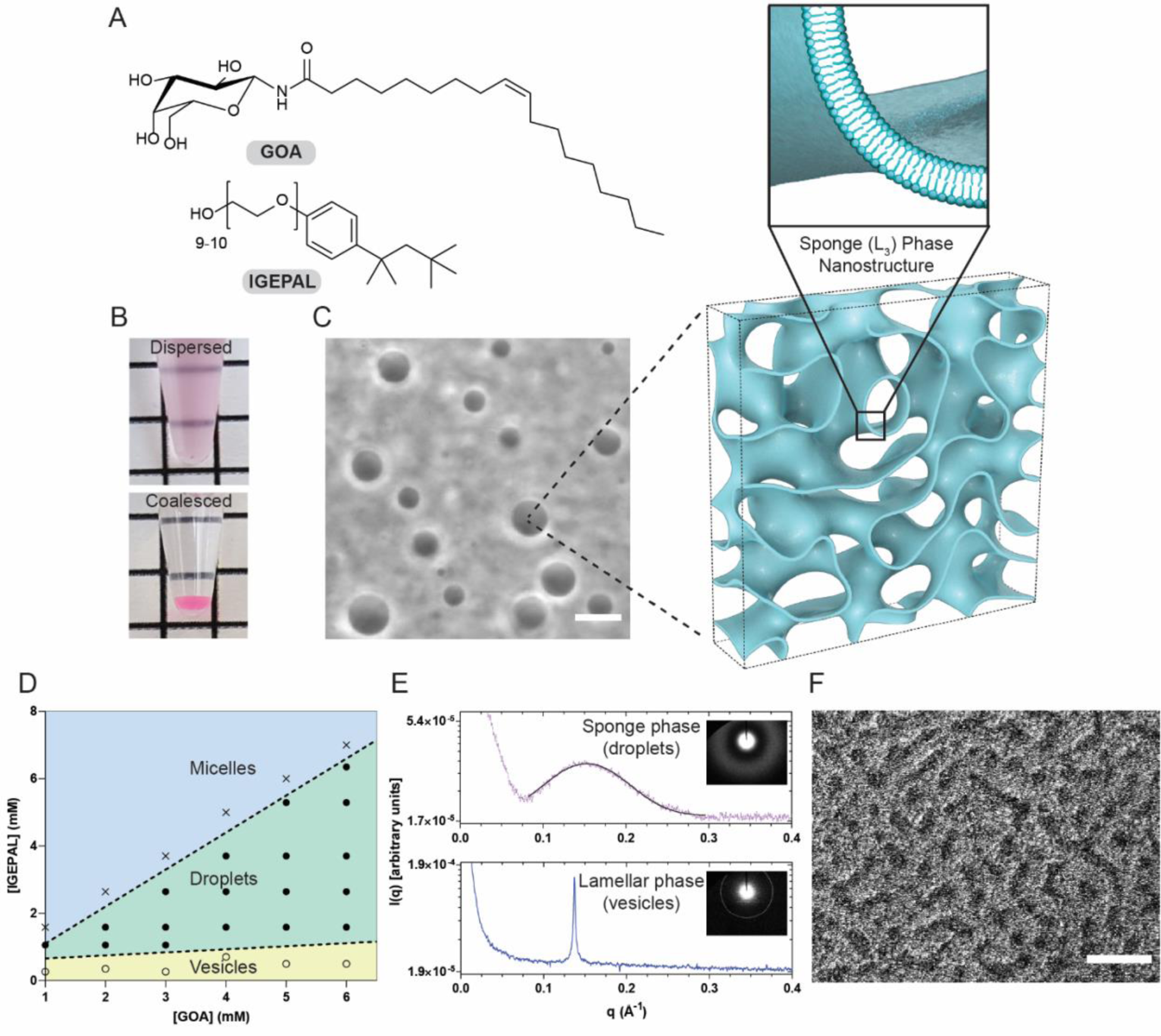
Physical characterization of the lipid sponge droplets. (*A*) Chemical structures of *N*-oleoyl β-D-galactopyranosylamine (GOA) and octylphenoxypolyethoxyethanol (IGEPAL). (*B*) A dispersed sample of droplets merges into a single coalesced phase at the bottom of the tube under gravity. A small quantity of the water soluble dye Rhodamine B was added to the dispersion to demarcate the coalesced phase. (*C*) Phase-contrast image of a typical droplet dispersion and illustration of the droplets’ porous, bilayer-rich nanostructure. Illustration was based on a computationally generated model of a bicontinuous structure (50). Scale bar: 10 µm. (*D*) Phase diagram showing the relative molar compositions of GOA and IGEPAL in 1X PBS over which vesicles (*yellow*), droplets (*green*), and micelles (*blue*) are obtained. (*E*) Top: synchrotron small-angle X-ray scattering (SAXS) intensity profile (dotted) from a droplet dispersion (4:3 GOA:IGEPAL by molar ratio). The solid black line represents a Gaussian fit to the data to obtain the position of maximum. Bottom: synchrotron SAXS intensity profile from a dispersion of GOA vesicles. The inset images show the SAXS patterns. (*F*) Freeze-fracture cryogenic scanning electron microscopy (cryo-SEM) image corresponding to sponge phase morphology. Scale bar: 1 µm.

## Results and discussion

### Formation and physical characterization of lipid sponge droplets

We synthesized *N*-oleoyl β-D-galactopyranosylamine (GOA, **Fig. 1*A***) as previously described (25). When a thin lipid film of GOA deposited on the walls of a glass vial is hydrated with a solution containing the non-ionic detergent IGEPAL (**Fig. 1*A***), a uniformly turbid dispersion is produced. On centrifugation or long standing, the dispersion coalesces into an optically isotropic single dense phase at the bottom of the tube (**Fig. 1*C***). By phase contrast microscopy, the dispersion was found to consist of a polydisperse population of micron-sized spherical droplets with distinct optical contrast compared to the surrounding medium (**Fig. 1*C***). The spherical shape suggested to us that the droplets are liquid in nature. The coalesced phase can be resuspended by gentle agitation or vortexing. We used high-performance liquid chromatography and mass spectrometry (HPLC-MS) to determine the compositions of the supernatant and the coalesced phases and found that both amphiphiles had partitioned almost entirely (∼99%) into the coalesced phase (***SI Appendix*, Fig. S1**). When thermogravimetric analysis (TGA) was carried out on the coalesced phase (obtained from 4:3 GOA:IGEPAL by molar ratio), we observed a gradual loss of about 67% mass over the range 30-100 °C (***SI Appendix*, Fig. S2**). This lost mass is likely originating from water present in the coalesced phase. We acquired the fluorescence emission spectrum of the solvatochromic dye Laurdan incorporated in the droplets and obtained a maximum around 490 nm (***SI Appendix*, Fig. S3**). This data suggests that the droplets have a water-accessible environment where the dye can undergo dipolar relaxation typical of fluid lipid membranes. In comparison, in oil-in-water droplet systems such as oleic acid, Laurdan exhibits a maximum peak at 440 nm. We constructed a phase diagram by varying the relative proportions of GOA and IGEPAL and found that droplet formation takes place at molar ratios of approximately 1:1 to 1:4 (IGEPAL:GOA) occupying a large portion of the phase space (**Fig. 1*D***). Outside this region, micellar solutions (at higher relative IGEPAL concentrations) or lamellar vesicles (at lower relative IGEPAL concentrations) are obtained.

We carried out small-angle X-ray scattering (SAXS) investigation on droplet dispersions using a synchrotron radiation source. When the intensity [*I*(*q*)] profile from a 4:3 GOA:IGEPAL droplet dispersion was plotted against the scattering vector (*q*), a single broad peak was obtained (**Fig. 1*E***). The peak is fitted to a Gaussian distribution and a maximum is obtained at *q*_max_ = 0.153±0.0002 Å^-1^. For various compositions of the droplets, the intensity profile showed a decay behavior as *I*(*q*) ≈ *q*^*-2*^ consistent with a local bilayer structure (***SI Appendix*, Fig. S4**) (26). Such features are typical of a lipidic sponge (L_3_) phase (**Fig. 1*C***) similar to that reported with various surfactant systems (24, 26). The characteristic length (*d* = 2π/*q*_*max*_) for the sponge phase (from 4:3 GOA:IGEPAL) was calculated to be 4.11 nm. This length can be interpreted as the average channel diameter of the sponge phase (24). We found that the nature of the peak and the position of the maxima of the SAXS profile remains nearly unchanged in the coalesced phase, suggesting that the structural characteristics are retained (***SI Appendix*, Fig. S5**). In comparison, SAXS on a dispersion of GOA vesicles gave rise to an intensity profile consisting of a sharp peak (**Fig. 1*E***) consistent with a lamellar (L_α_) phase (27).

We further characterized the droplets by cryogenic transmission electron microscopy (Cryo-TEM) and found that they lack any well-ordered internal structure, consistent with the characteristics of a sponge phase (***SI Appendix*, Fig. S6*A***) (28). In contrast, Cryo-TEM analysis of a sample of GOA vesicles showed distinct membrane bound structures of various lamellarities (***SI Appendix*, Fig. S6*B***). We imaged the droplets by freeze-fracture cryogenic scanning electron microscopy (Cryo-SEM) and observed a porous morphology (**Fig. 1*F***) corresponding to a three-dimensional network of lipid bilayers (19, 22, 29). In addition to IGEPAL, we found that GOA forms droplets with other nonionic octylphenoxypolyethoxyethanol detergents such as Triton X-114 and Triton X-100 (***SI Appendix*, Fig. S7**). Single chain galactolipids with shorter unsaturated fatty acid chains (25) such as GPOA and GMOA were also found to form droplets with the same class of non-ionic surfactants (***SI Appendix*, Fig. S7**). Previously, it has been proposed that transformations of lamellar to sponge phases are mediated by bending of bilayers followed by formation of interconnections between them (18). GOA and similar galactolipids alone form lamellar phases, which are composed of locally flat stiff bilayer membranes. We speculate that octylphenoxypolyethoxyethanol surfactants, with their flat aromatic ring and short aliphatic segment, insert between the GOA molecules and induce a negative curvature. Additionally, the detergents may make the bilayers more fluid by lowering the bending rigidity. These combined effects would allow adjacent bilayers to fuse and form a three-dimensional sponge-like network.

We found that stable droplets can be formed over a wide range of pH, ionic strength, buffer composition, and dilution. We observed droplet formation over the pH range 2-10, high ionic strength (up to 1 M NaCl), high concentrations of Mg^2+^ (100 mM) and Ca^2+^ (10 mM) (***SI Appendix*, Fig. S8*A*-*B***). The droplets also formed in complex media like sea water, cell culture media, and cell-free transcription/translation (TX-TL) reactions (***SI Appendix*, Fig. S8*C*-*E***). Due to lack of an ionizable head group, glycolipid sponge droplets assemble more robustly compared to other droplet systems based on single chain amphiphiles, such as fatty acids (30), which can exist only within a relatively narrow range of pH conditions and ionic environments.

### Partitioning of small molecules and fluorescent dyes

By virtue of their extensive bilayer network and high water content, lipid sponge droplets can provide a unique milieu for harboring molecules spanning from hydrophobic to hydrophilic in nature. At first, we studied the partitioning behavior of a variety of molecules into the sponge phase using HPLC-MS. We measured the concentration of an analyte in the supernatant and compared it with the total concentration to estimate the percentage of partitioning into the sponge phase (***SI Appendix*, Table S1**). Polar lipid molecules such as cholesterol and 1-palmitoyl-2-oleoyl-glycero-3-phosphocholine (POPC) were 100% partitioned. Water soluble fluorescent dye Rhodamine B was 81.4% partitioned while the more polar dye Sulforhodamine B was 47.3% partitioned. Next we used fluorescence microscopy to visualize the partitioning behavior of several lipophilic, and water soluble fluorescent molecules (**Fig. 2*A, SI Appendix*, Fig. S9*A***). Highly sulfonated dyes such as Alexa Fluor 488, and sulfo-Cy5-azide showed little or no enrichment (**Fig. 2*A*, vii-viii**). This suggests that dyes like Alexa Fluor 488 can be used as a fluorescent label for macromolecules to study their localization into the droplets in an unambiguous manner. Non-sulfonated xanthene dyes such as fluorescein and 4’,5’-dibromofluorescein showed different degrees of partitioning depending on pH and hence their ionization state (***SI Appendix*, Fig. S9*B*-*C***). For instance, at lower pH (∼6.0) the phenolic -OH of fluorescein remains mostly protonated, resulting in greater partitioning (43.7%) to the sponge phase. At higher pH (∼8.0), the phenolic -OH is ionized, leading to significantly less partitioning (13.2%) to the sponge phase. Based on the above studies, we conclude that overall charge, hydrophobicity/hydrophilicity, and structure of the molecules play an important role in determining the partitioning behavior.

**Fig. 2.**
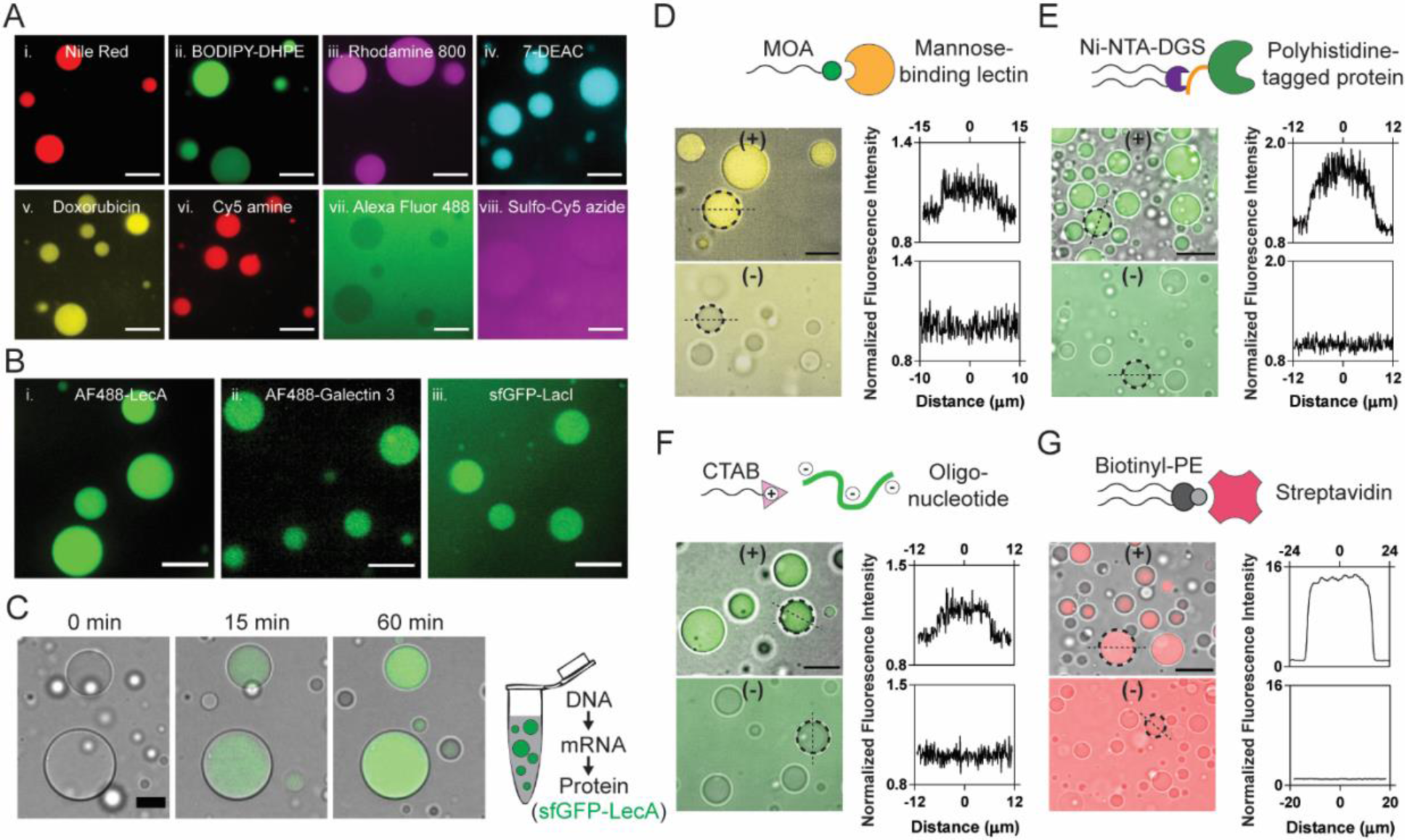
Partitioning of various molecules into lipid sponge droplets. (*A*) Partitioning of i) Nile Red ii) BODIPY FL-DHPE iii) Rhodamine 800 iv) 7-diethylaminocoumarin v) Doxorubicin vi) Cy5-amine vii) Alexa Fluor 488 and viii) sulfo-Cy5-azide into lipid sponge droplets. Scale bars: 10 µm. (*B*) Partitioning of galactophilic proteins i) Alexa Fluor 488-LecA ii) Alexa Fluor 488-Galectin 3 and iii) sfGFP-LacI into lipid sponge droplets. Scale bars: 10 µm. (*C*) Time lapse images showing gradual incorporation of sfGFP-LecA expressed in PURE system into lipid sponge droplets. Scale bar: 15 µm. (*D*) Selective partitioning of FITC-Concanavalin A into droplets doped with *N*-oleoyl β-D-mannosylamine (MOA). Scale bar: 15 µm. (*E*) Selective partitioning of sfGFP-His_6_ into droplets doped with Ni-NTA-DGS. Scale bar: 20 µm. (*F*) Selective partitioning of a FAM-labeled DNA oligonucleotide into droplets doped with CTAB. Scale bar: 15 µm. (*G*) Selective partitioning of Alexa Fluor 568-labeled streptavidin into droplets doped with biotinyl-PE. Scale bar: 35 µm. In (*D*)-(*G*), binding (doped) droplets are represented by (+) while non-binding droplets are represented by (-). Plots of the fluorescence intensity profile along a straight line drawn through the center of the marked droplets (broken circle) are shown on the right. On the *x*-axes, 0 corresponds to the center of the droplets.

### Partitioning of galactophilic proteins

Galactose and its derivatives play a crucial role in biology as affinity or recognition groups for protein binding on the surfaces of cells or organelles. GOA offers ample β-galactopyranosyl moieties throughout the bulk of the sponge droplets. We hypothesized that proteins with affinity for galactose residues would spontaneously partition into the droplets. We chose to test the well-studied galactophilic lectin LecA (PA-IL) from the pathogenic bacterium *Pseudomonas aeruginosa*, where LecA promotes biofilm formation via cross-linking between bacterial cell-surface galactose residues (31). We added Alexa Fluor 488-labeled LecA (1 µM final concentration) to droplets (formed from 3 mM GOA and 1.8 mM IGEPAL) and observed a high extent of partitioning into the droplets (**Fig. 2*B*-i**). Using fluorometric analysis of the supernatant, we estimated that 97.3% of the protein partitioned into the droplets. We carried out control experiments where isopropyl β-D-1-thiogalactopyranoside (IPTG) or phenyl β-D-galactopyranoside were added as competing ligands for LecA and significantly less partitioning of the Alexa Fluor 488-labeled LecA was observed by microscopy, thus supporting our hypothesis that sequestration was due to a specific interaction of the lectin with GOA (***SI Appendix*, Fig. S10**). Next we tested the galactophilic lectin galectin 3, which is known to bind primarily to β-galactoside residues and plays an important role in cell adhesion, macrophage activation, and apoptosis (32). As expected, we observed highly efficient partitioning of Alexa Fluor 488-labeled galectin-3 into the droplets (**Fig. 2*B*-ii**). Apart from galactophilic lectins, we found that a GFP fusion of the *Escherichia coli lac* operon repressor protein (sfGFP-LacI) spontaneously partitions into the droplets (**Fig. 2*B*-iii**). LacI is known to bind to galactopyranoside effector ligands primarily via hydroxyl groups (O2, O3, O4, O6) of the galactose moiety on the latter. In addition, apolar substituents (if any) on the C1 (anomeric) position of galactose group interact with a hydrophobic surface in the binding site, thereby increasing the binding affinity (33). We reason that the presence of the galactose group and hydrophobic oleoyl chain on GOA contribute to its binding to sfGFP-LacI.

### Cell-free protein synthesis in the presence of droplets

To evaluate the compatibility of droplets and their sequestration properties with complex biochemical reactions, we formed droplets in a recombinant TX-TL system (PURExpress). The PURE TX-TL system consists of almost 30 enzymes, 56 tRNAs, ribosomes, and a highly optimized mixture of precursors and buffer components (34). Even when droplets were present at high densities (reaction contained 12 mM GOA and 6.7 mM IGEPAL), efficient protein synthesis took place and we observed gradual sequestration of galactophilic proteins during their synthesis (**Fig. 2*C*, *SI Appendix*, Fig. S11, *SI Appendix*, Video S1**). While LecA and LacI fused with sfGFP spontaneously partitioned into the droplets upon synthesis, sfGFP alone did not (**Fig. 2*C*, *SI Appendix*, Fig. S11**). Cell-free protein synthesis in the presence of droplets shows that droplets are highly biocompatible and will likely support many different biochemical reactions. Further, we can utilize this methodology as a rapid route to introduce functional proteins into the droplets to carry out specific biochemical transformations.

### Programming partitioning behavior

Living cells and sub-cellular organelles contain specific receptors to recognize macromolecular binding partners or signaling molecules. For example, lysosomal enzymes synthesized in the rough ER are modified with mannose-6-phosphate for targeting them to lysosomes (35). Since our droplets are composed of amphiphilic species, we asked if such targeting mechanisms could be mimicked by doping the droplets with amphiphiles bearing affinity handles. At first, we doped the droplets with 11.3 mol% *N*-oleoyl β-D-mannopyranosylamine (MOA) and added fluorescein isothiocyanate (FITC) labeled lectin Concanavalin A, which has specificity for terminal mannopyranosyl residues. As expected, we observed that the droplets doped with MOA had a 1.14-fold higher fluorescence signal compared to the background while those without did not show increased fluorescence (**Fig. 2*D***). Encouraged by this result, we doped the droplets with 1.6 mol% 1,2-dioleoyl-*sn*-glycero-3-[(*N*-(5-amino-1-carboxypentyl) iminodiacetic acid) succinimidyl] (nickel salt) (Ni-NTA-DGS) – a headgroup modified phospholipid that binds to polyhistidine-tagged proteins with high affinity and specificity. When we added sfGFP-His_6_, we observed highly efficient localization using confocal microscopy. In absence of Ni-NTA-DGS, no localization of sfGFP-His_6_ was observed (**Fig. 2*E***). To further quantify how much sfGFP-His_6_ partitioned into Ni-NTA-DGS-doped droplets, we analyzed droplet supernatants fluorometrically. Droplets sequestered more than 90% of the protein when we titrated sfGFP-His_6_ between 1 nM and 10 µM (***SI Appendix*, Fig. S12**). At 10 µM total sfGFP-His_6_ concentration, we estimate the protein concentration to be almost 1 mM in the droplet phase, based on droplet formation experiments at comparable GOA and IGEPAL concentrations in large volumes, where the droplets phase occupied approximately 1% of the total volume. Our calculations indicate that His-tagged proteins in the droplet phase can therefore easily reach the concentrations of even the most abundant proteins in cells (for instance, the translation elongation factor EF-Tu is present at approximately 100 µM in *E. coli*) (17). The capacity of droplets for sequestering His-tagged protein obviously depends on the density of droplets and the amount of Ni-NTA-DGS, which explains why partitioning saturates as the protein concentration approaches the concentration of Ni-NTA-DGS (***SI Appendix*, Fig. S12**). Next, we doped the droplets with 10 mol% of the cationic amphiphile cetyltrimethylammonium bromide (CTAB) and found that it facilitated the sequestration of a fluorescently labeled oligonucleotide (5’-FAM dN_20_) (**Fig. 2*F***). We measured the fluorescence signal in the droplets to be 1.2-fold higher than the background when 2 µM 5’-FAM dN_20_ was added to CTAB-doped droplets, while no increase was observed in non-doped droplets. Doping with a biotinylated phospholipid (biotinyl-PE) allowed us to recruit fluorescently labeled Streptavidin into the sponge phase, where the protein was concentrated more than 10-fold from solution (**Fig 2*G***). Recruitment of streptavidin suggests that additional biotinylated molecules may be co-sequestered when bound to streptavidin tetramers.

### Mobility of molecular cargo in lipid sponge droplets

For reactions to occur in a synthetic organelle model, it is important that molecules are mobile within the compartment. As mentioned earlier, the spherical shape suggests that the droplets are liquid in nature. Additionally, we frequently observe fusion events between individual droplets that coalesce into a bigger droplet (***SI Appendix*, Video S2**). To quantify the mobility of molecules inside droplets, we performed fluorescence recovery after photobleaching (FRAP) experiments (***SI Appendix*, Fig. S13**). To study the mobility of hydrophobic molecules, we bleached the center of a droplet containing a fluorescently labeled phospholipid and the half-maximal recovery of fluorescence was observed within 6.8 s (t_1/2_). To study the mobility of soluble proteins, we carried out FRAP on Ni-NTA-DGS-doped droplets binding a His_6_-tagged fluorescent protein and measured a t_1/2_ of 16 s. As controls, we compared protein diffusion in droplets to the mobility of the protein in aqueous water-in-oil emulsion droplets and bound to a Ni-NTA gel matrix, where, respectively, fluorescence recovery was much more rapid or did not occur over the duration of the experiment (***SI Appendix*, Fig. S13**). The FRAP results indicate that molecular cargo within the droplet phase diffuses slower than in a dilute aqueous solution but they are sufficiently mobile to allow dynamic molecular interactions to occur.

### Reconstitution of functional transmembrane proteins

Membrane proteins are encoded by a large proportion (∼30%) of the genome and they fulfill essential cellular functions (20). However, they are challenging to employ in systems built from the bottom up, limiting the range of biochemical transformations that can be implemented in cell-free reactions. Since the droplets are composed of biocompatible lipid bilayers, we were interested in testing if spontaneous reconstitution of transmembrane proteins was possible. As a proof of concept demonstration, we chose commercially available cytochrome c oxidase (CcO) solubilized in *n*-dodecyl-*β*-D-maltoside (DDM) as a model integral membrane protein. CcO is a large transmembrane protein complex found in the inner mitochondrial membrane and it is the terminal enzyme of the respiratory electron transport chain. Initially, we exchanged DDM with IGEPAL using spin filtration. Next, we prepared the droplets by hydrating a GOA film in the presence of IGEPAL-solubilized CcO. When ferrocytochrome c was added to the droplets, we observed a steady decrease in absorbance at 550 nm indicating the enzyme-catalyzed formation of ferricytochrome c (**Fig. 3*A***). In comparison, when the activity was measured with IGEPAL-solubilized CcO only, a two-fold lower reaction rate was observed. The bilayer membrane environment within the droplets likely allowed proper folding and membrane insertion of CcO in comparison to solubilization in IGEPAL only. Finally, for direct observation, we labeled CcO with Alexa Fluor 488, and observed spontaneous partitioning into the droplets (**Fig. 3*A***).

**Fig. 3.**
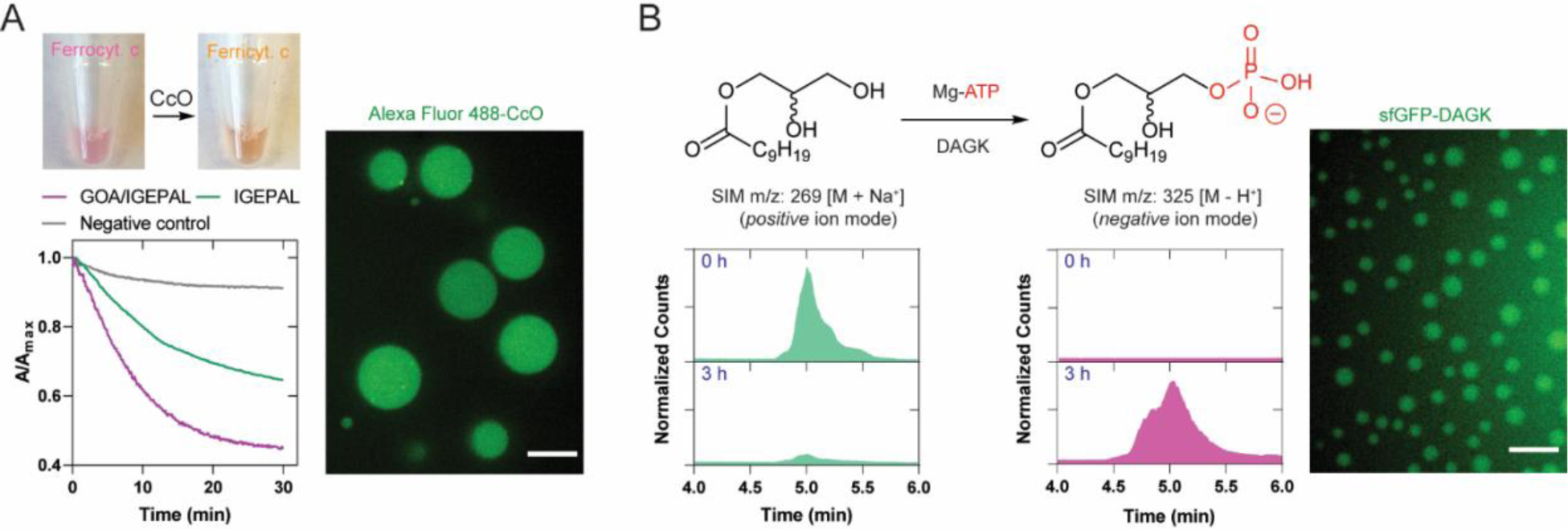
Reconstitution of functional transmembrane proteins in lipid sponge droplets. (*A*) Cytochrome c oxidase (CcO) reaction. Addition of ferrocytochrome c to a droplet dispersion containing CcO leads to a gradual decrease in the absorbance at 550 nm indicating the formation of ferricytochrome c. Alexa Fluor 488-labeled CcO partitions into the droplets. Scale bar: 10 µm. (*B*) *E. coli* diacylglycerol kinase (DAGK) reaction in droplets. DAGK is expressed in PURE system in the presence of droplets containing the substrate 1-decanoyl-*rac*-glycerol. Selective ion monitoring (SIM) chromatogram shows transformation to the phosphorylated product. Similarly expressed fluorescent fusion protein sfGFP-DAGK partitions into the droplets. Scale bar: 5 µm.

Membrane proteins of the endomembrane system participate in a variety of transformations of lipid substrates. The ER contains the biosynthetic machinery for the production of phospholipids and steroids (36). Comparably packed with membrane bilayer networks and with their ability to sequester membrane proteins and hydrophobic molecules, sponge droplets may offer an excellent environment for reactions involving lipid substrates. To test our hypothesis, we expressed bacterial (*E. coli*) diacylglycerol kinase (DAGK), a 13.2 kDa protein containing three transmembrane helices, in a PURE TX-TL reaction in the presence of droplets. DAGK catalyzes the phosphorylation of diacylglycerols and monoacylglycerols. The droplets were doped with 9.1 mol% 1-decanoyl-*rac*-glycerol, a substrate for DAGK, and additional ATP and MgCl_2_ were added in the reaction mixture to support phosphorylation activity. After 3 h, the expected product decanoyl lysophosphatidate could be detected in nearly quantitative conversion as evidenced by the mass spectrometric signals (**Fig. 3*B***). To visualize the partitioning of the protein into the droplets, we expressed sfGFP-fused DAGK in the TX-TL system and observed fluorescence signal in the droplets indicating localization of the expressed protein (**Fig. 3*B***). Our results demonstrate that droplets can facilitate an important enzymatic step in the biosynthesis of phospholipids. In the future, lipid sponge droplets, with their high membrane density and large lipid-aqueous interface, could provide a platform for reconstituting more complex biochemical pathways involving multiple transmembrane proteins. In particular, we believe there is potential for the reconstitution of lipid biosynthesis pathways to emulate and characterize functions of the smooth ER.

### Rate enhancement of enzymatic reactions in the droplets

Living cells use organelles to co-localize molecules to facilitate reactions and minimize unwanted side reactions. Using compartmentalization to enhance biochemical reaction rates in living cells has been demonstrated by synthetic biologists and metabolic engineers. For instance, confinement of enzymatic pathways and key intermediates in mitochondria of engineered yeast has been shown to increase metabolite titer significantly as compared to cytoplasmic pathways (37). Similar strategies have been employed to enhance the yield of fatty acid-derived metabolites in yeast peroxisomes (38). We asked if colocalization of an enzyme and its substrate in droplets would enhance the rate of a biochemical reaction. First, we bound His_6_-tagged cathepsin K, a lysosomal cysteine protease, within Ni-NTA-DGS doped droplets (**Fig. 4*A***). Next, we added a fluorogenic dipeptide substrate benzyloxycarbonyl-L-leucyl-L-arginine 7-amido-4-methylcoumarin (Z-LR-AMC), which gets sequestered within droplets by virtue of its partial hydrophobic nature (**Fig. 4*A***). We monitored the progress of the cathepsin K catalyzed hydrolysis reaction by measuring the steadily increasing fluorescence signal from the by-product 7-amino-4-methylcoumarin (AMC) (**Fig. 4*B***). We observed that the progress of reaction was markedly higher when compared to the bulk phase reaction in the absence of droplets. We performed additional control reactions where we (i) omitted Ni-NTA-DGS from the droplets (non-binding droplets), and (ii) used IGEPAL only, and in each case, we found that the increase in AMC fluorescence signal was lower than the condition where both enzyme and substrate were co-localized in the droplets. We further corroborated and quantified these results by measuring the amount of AMC generated using HPLC-MS based on standard calibration curves (***SI Appendix*, Fig. S14**). We estimated that after 6 hours, 84.7% of Z-LR-AMC was hydrolyzed in the binding droplets (**Fig. 4*C***). In the presence of non-binding droplets or IGEPAL alone, 25.7% and 18.6% of Z-LR-AMC was hydrolyzed respectively. Only 2.5% of Z-LR-AMC was hydrolyzed in the bulk condition. A likely explanation for the higher yield of product in the non-binding droplets and IGEPAL only conditions compared to the bulk without additives is that the amphiphiles facilitated dispersal of the substrate in the reaction media. Nonetheless, the product yield is enhanced greatly by programming the droplets to sequester cathepsin K and by concentrating and co-localizing enzyme and substrate. Previously enhanced reaction rates have been observed by concentrating biomolecules in complex coacervates (4, 39, 40). We believe that the lipid sponge droplets will offer a general programmable scaffold for studying the effects of colocalization and confinement on enzymatic reactions.

**Fig. 4.**
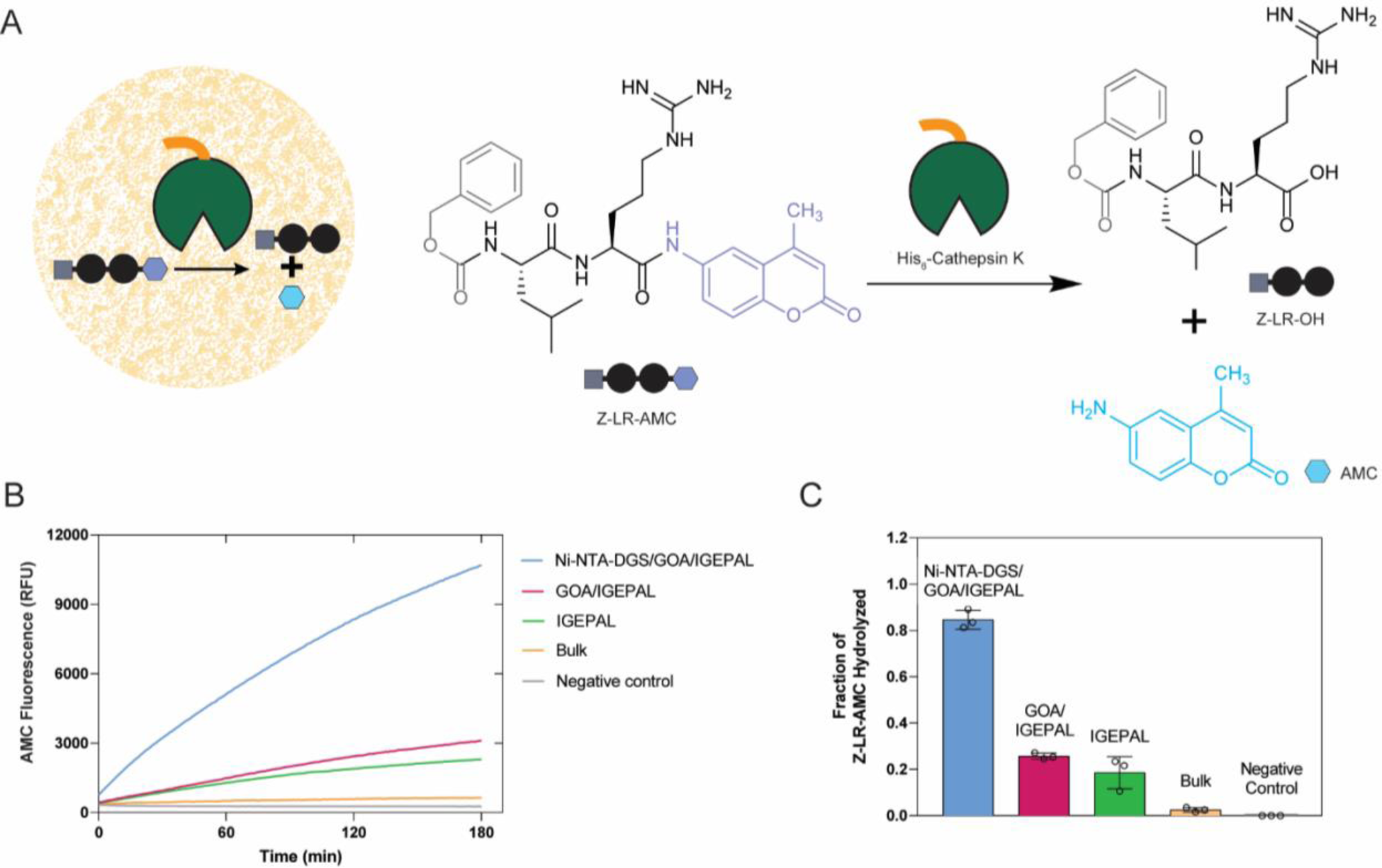
Rate enhancement of a model biochemical reaction due to co-localization of an enzyme and its substrate. (*A*) Reaction scheme showing hydrolysis of a fluorogenic dipeptide Z-LR-AMC catalyzed by His_6_-Cathepsin K. (*B*) Progress of hydrolysis of Z-LR-AMC in droplets and corresponding controls followed by fluorometry. (*C*) Quantification of the extent of hydrolysis of Z-LR-AMC in droplets and corresponding controls by HPLC-MS.

### Light-controlled protein encapsulation

In living cells, proteins often shuttle dynamically between different subcellular locations in response to changes in the environment. For example, transcription factors shuttle between the nucleus and the cytoplasm, and cells dynamically alter the number of signaling receptors in the plasma membrane by endocytosis (41). Furthermore, cells regulate the availability of signaling proteins by forming and dissolving membraneless compartments such as stress granules in response to environmental factors (1). Sequestration of proteins into intracellular phase separated droplets has also been engineered to be responsive to light stimuli (42). We aimed to engineer light-controlled and reversible protein sequestration into lipid sponge droplets. The photoreceptor phytochrome B (PhyB) from *Arabidopsis thaliana* switches between a binding and a non-binding conformation upon exposure to red (660 nm) and far-red (740 nm) wavelengths respectively (***SI Appendix*, Fig. S15**). In its binding conformation, PhyB binds to phytochrome interacting factors (PIFs) that can be fused to cargo proteins (43). We used this system to control the capture and release of proteins by droplets with light. PhyB was produced with a biotin affinity handle, bound to Streptavidin and encapsulated in droplets doped with biotinyl-PE (**Fig. 5*A***). PhyB-loaded droplets encapsulated PIF-sfGFP-ssrA after exposure to daylight or 660 nm light. The protein was not enriched in droplets in 740 nm light or in the absence of PhyB (**Fig. 5*B*, *SI Appendix*, Fig. S16**). Alternating exposure of droplets with red and far-red light reversibly switched the localization of PIF-sfGFP-ssrA between droplets and solution. Release and capture of the protein was rapid. Droplets lost 90% of GFP fluorescence within 2-3 min after switching illumination to 740 nm far-red light, and fluorescence signal from the solution in the vicinity of droplets increased immediately. When droplet samples were illuminated with 660 nm light, fluorescence signal from the solution decreased and fluorescence of droplets began to increase. Fluorescence of PIF-sfGFP-ssrA was observed to spread from the edges into the interior of droplets. In large droplets with diameters of more than 40 µm it took more than 30 min to reach a homogeneous intensity of GFP from droplet edge to interior (**Fig. 5*B*, *SI Appendix*, Video S3**). The observed differences in the release and binding speeds can be explained by the differences in PIF-tagged protein concentration under the different illumination conditions in droplets and solution respectively. Sequestration into droplets is slower than release because the protein is at a relatively low concentration in solution, and sequestration into the droplets only occurs after an encounter of a PIF-tagged protein in solution and a droplet-bound PhyB. The observed fast release kinetics will be useful to rapidly make proteins available for reaction in solution. Light controlled localization changes were readily reversible and could be observed for more than 6 h, which highlights the advantage of a porous material such as lipid sponge droplets to control localization of molecules. In contrast, separation of molecules in nested vesicle-in-vesicle assemblies is typically irreversible (44).

**Fig. 5.**
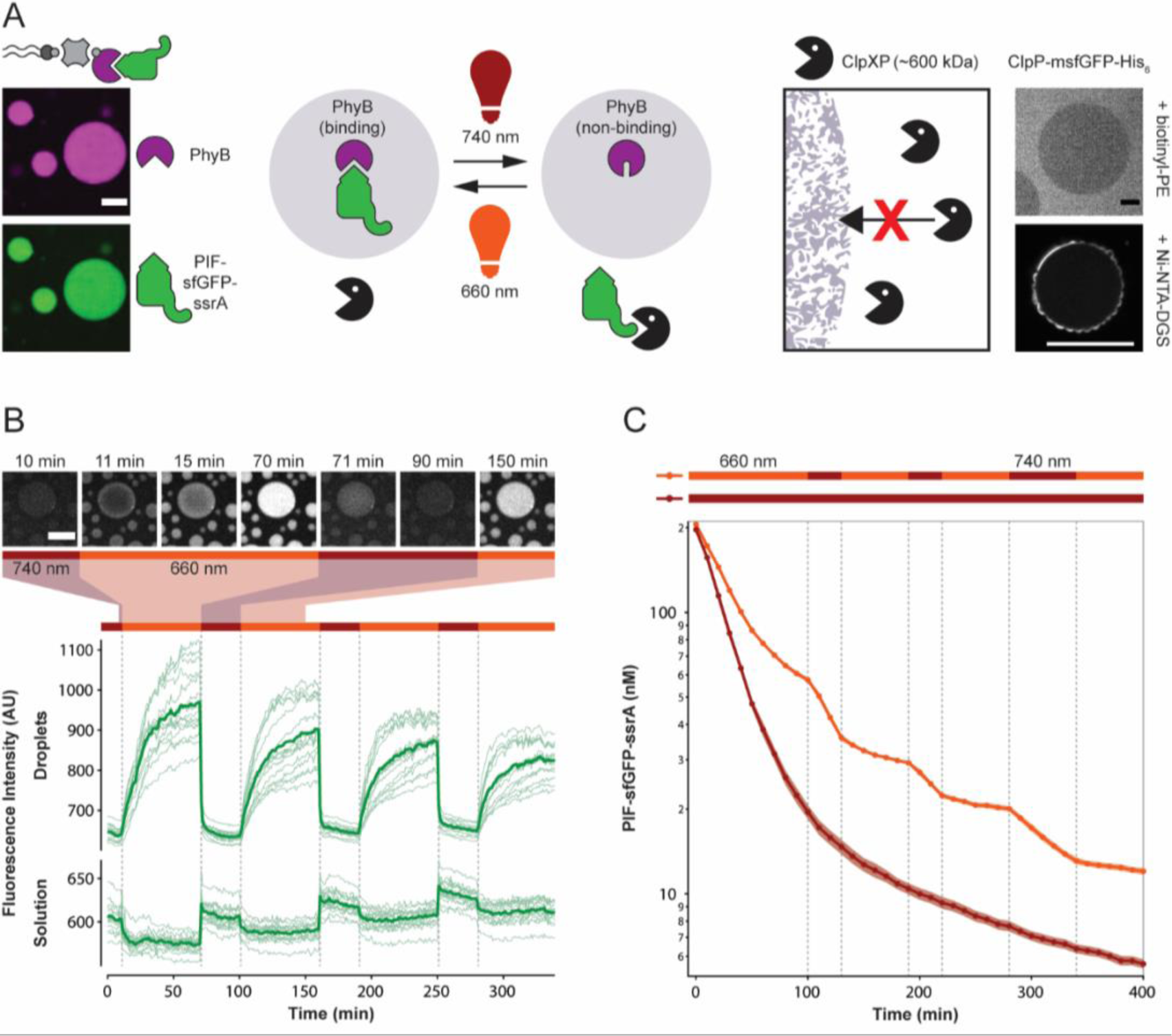
Light-dependent protein capture and release to dynamically control degradation rates. (*A*) Design of the light-controlled protein degradation reaction. Left: Biotin-Streptavidin-mediated binding of light-responsive phytochrome B (PhyB) and capture of the protease substrate PIF-sfGFP-ssrA in 660 nm light. Right: Exclusion of ClpXP protease from lipid sponge droplets. ClpP-msfGFP-His_6_ is excluded from droplets doped with biotinyl-PE and is unable to enter the droplets when targeted inside via Ni-NTA-DGS. (*B*) Light-controlled release and capture of PIF-sfGFP-ssrA by PhyB-loaded droplets. Traces show fluorescence intensities of droplets and solution over time in response to changing illumination conditions. Bars above indicate 660 nm and 740 nm conditions. Intensities were extracted from a total of 15 droplets in 5 time-lapse movies (thin lines) and averaged (bold lines). Selected images from a time-lapse movie are shown on top. (*C*) Light-controlled switching between protection and degradation of PIF-sfGFP-ssrA by ClpXP. Degradation of PIF-sfGFP-ssrA was monitored by reading the fluorescence of the reaction every 10 min. Reactions were performed in triplicates. Shown is the average (bold) and the standard deviation (shaded areas). Scale bars: 20 µm.

### Light-controlled sequestration in droplets reversibly protects from degradation

Cells utilize a “nucleolar detention” strategy to regulate molecular networks by capturing and immobilizing essential cellular factors within the nucleolus away from their effector molecules (45). We reasoned that light-controlled protein release and capture by droplets should allow us to control reactions in a similar manner. In contrast to the previous reactions that we engineered to occur within droplets, we focused on a reaction happening in solution, outside of droplets: ClpXP mediated protein degradation. ClpXP (from *E. coli*) is a large protein complex of approximately 600 kDa that consists of 6 ClpX and 14 ClpP subunits. ClpXP degrades ssrA-tagged protein substrates in a highly specific, processive and ATP-dependent manner and has a K_M_ of about 500 nM for ssrA-tagged sfGFP (46). Since this protein complex has a cylindrical shape of 14.5 nm length and 11 nm diameter (47), which are significantly larger than the average channel diameter of 4.1 nm, we expect it to be excluded from the droplets. Indeed we observe that ClpP-msfGFP-His_6_ is excluded from biotinyl-PE-doped droplets and cannot enter the droplets even when targeted inside using Ni-NTA-DGS (**Fig. 5*A*, *SI Appendix*, Fig. S17**). Light-controlled changes in PIF-sfGFP-ssrA concentration in solution should therefore affect the rate of proteolytic degradation of the protein by ClpXP. As predicted, we found that in 660 nm illumination conditions degradation of PIF-sfGFP-ssrA was substantially slower than in 740 nm illumination conditions in the presence of PhyB-loaded droplets. We observed differences in degradation rate by fluorometric measurements of the reaction over time (**Fig. 5*C*, *SI Appendix*, Fig. S18**). We further corroborated these data by microscopy of reaction samples, and immunoblotting (***SI Appendix*, Figs. S19-20**). The light-dependent behavior allowed us to dynamically control the degradation rate by changing the illumination wavelength. Switching illumination between 660 nm and 740 nm led to pronounced and immediate changes in degradation rate (**Fig. 5*C***). Protection from ClpXP-mediated degradation required the presence of PhyB-loaded droplets. The Streptavidin-PhyB complex alone, without droplets, slowed degradation only marginally. Also, in the presence of droplets without PhyB, no protection or responses to wavelength changes were observed. In the absence of ClpXP, GFP signal decreased only marginally (***SI Appendix*, Fig. S18**). As shown here for light-switchable protection of a sequestered protein from degradation in solution, light-responsive droplets will enable precise spatiotemporal control of reactions both in solution and in the droplet phase. In principle, light-controlled sequestration into droplets can be applied to any soluble protein offering much greater flexibility in reaction control than complex coacervate systems that have been previously engineered to reversibly assemble and disassemble in response to enzyme activity (48) or temperature (49).

## Conclusion

In summary, we describe the application of lipid sponge droplets as mimics of membrane-rich organelles and subcellular structures. With their nanoporous structure offering abundant hydrophobic-hydrophilic interfaces, the droplets spontaneously sequester molecules from the surrounding medium based on their physico-chemical properties. Further, the droplets are programmable in a modular fashion using amphiphiles with affinity handles that specifically target tagged proteins into the sponge phase at high concentrations. This is a significant advantage over vesicle-based artificial organelle models, which suffer from the limitation that they have no mechanism for efficient and spontaneous internal sequestration of macromolecules and solutes. Being composed of biocompatible building blocks, the lipid sponge droplets support a wide range of enzymatic reactions catalyzed by both transmembrane and soluble proteins and involving both water soluble and lipophilic substrates. Furthermore, we showed that a combination of colocalization, increased concentration, and the molecularly crowded environment can enhance enzymatic reactions in the droplets. The open and liquid nature of the droplets allows exchange with the surrounding environment, as well as reversible, light-responsive control over protein localization and concentration, which is difficult to engineer in a vesicle-based artificial organelle. Due to their versatile, self-assembled, and programmable nature lipid sponge droplets will find application as organelle-mimics to spatially organize biochemical reactions within artificial cellular systems and in cell-free systems. Finally, alternative, non-vesicular lipid structures such as the one reported here could serve as additional models for protocellular evolution. Fatty acid vesicles, which are popular protocell models, have been shown to transform to coacervates under certain conditions (30). Interconversion or coexistence of such structures may have important implications in protocellular structural diversity, stability, and permeability and deserves further exploration.

## Materials and Methods

Full experimental procedures of droplet characterization and biochemical methods are described in the ***SI Appendix***. Briefly, to form droplets, a thin lipid film was deposited on the walls of a glass vial from a solution of GOA (and, if required, amphiphiles bearing affinity handles) in chloroform and methanol by evaporation of the solvents. The lipid film was then rehydrated in a solution containing IGEPAL within the droplet forming composition range (**Fig. 1D**) by careful vortex mixing until a cloudy dispersion formed. Proteins or other additives were either supplied in the rehydration solution or added later as indicated for each experiment.

## Acknowledgements

This material is based upon work supported by the National Science Foundation (CHE-1844346; Devaraj) and the US Department of Energy [Biomolecular Materials Program, Division of Materials Sciences and Engineering] (DE-SC0018086; Sinha). H.N. was supported by a Swiss National Science Foundation fellowship (P300PA_174346). R. J. B thanks the Human Frontier Science Program (HFSP) for his Cross-Disciplinary Fellowship. We thank Soenke Seifert (Advanced Photon Source, Argonne National Laboratory) for guidance and valuable suggestions with the Synchrotron SAXS experiments, Simpson Joseph and Xinying Shi (UCSD) for ClpX and ClpP proteins, Maximilian Hörner and Wilfried Weber (University of Freiburg) for PhyB and PIF-tag plasmids, Florian Stenger and Axel Voigt (TU Dresden) for 3D models of a bicontinuous structure, and Veronica Falconieri Hays for illustration of the lipid sponge phase. TEM work was performed at the UC Irvine Materials Research Institute (IMRI). We acknowledge the use of the UCSD Cryo-Electron Microscopy Facility, which is supported by NIH grants to Timothy S. Baker and a gift from the Agouron Institute to UCSD. This work made use of the BioCryo facility of Northwestern University’s NU*ANCE* Center, which has received support from the Soft and Hybrid Nanotechnology Experimental (SHyNE) Resource (NSF ECCS-1542205); the MRSEC program (NSF DMR-1720139) at the Materials Research Center; the International Institute for Nanotechnology (IIN); and the State of Illinois, through the IIN. It also made use of the CryoCluster equipment, which has received support from the MRI program (NSF DMR-1229693). We thank Sebastian Maerkl (EPFL) and Itay Budin (UCSD) for valuable comments on the manuscript.

## Additional information

Any *SI Appendix*, information, chemical compound information and source data are available in the online version of the paper. Correspondence and requests for materials should be addressed to N.K.D.

