## Supplementary Information for "Lipid Sponge Droplets as Programmable Synthetic Organelles"

**This PDF file includes:**

Materials and Methods
Figures S1 to S21
Tables S1 to S4
Legends for Movies S1 to S3
SI References

**Other supplementary materials for this manuscript include the following:**

Movies S1 to S3

### Materials and Methods

**HPLC-MS**. Solvent mixtures for chromatography are reported as volume/volume (v/v) ratios. HPLC analyses were carried out on an Eclipse Plus C8 analytical column with *Phase A*/*Phase B* gradients [*Phase A*: $H_2O$ with 0.1% formic acid; *Phase B*: MeOH with 0.1% formic acid]. HPLC purification was carried out on Zorbax SB-C18 semipreparative column with *Phase A*/*Phase B* gradients [*Phase A*: $H_2O$ with 0.1% formic acid; *Phase B*: MeOH with 0.1% formic acid]. Electrospray Ionization-Mass spectra (ESI-MS) were obtained on an Agilent 6230 Accurate-Mass Time of Flight (TOF) mass spectrometer.

**Microscopy**. Spinning-disk confocal microscopy images were acquired on a Yokagawa spinning-disk system (Yokagawa, Japan) built around an Axio Observer Z1 motorized inverted microscope (Carl Zeiss Microscopy GmbH, Germany) with a 63x, 1.40 NA oil immersion objective or 20x 0.8 NA objective to an ORCA-Flash4.0 V2 Digital CMOS camera (Hamamatsu, Japan) using ZEN Blue imaging software (Carl Zeiss Microscopy GmbH, Germany). The fluorophores were excited with diode lasers (405 nm-20 mW, 488 nm-30 mW, 561 nm-20 mW, and 638 nm-75 mW). A condenser/objective with a phase stop of Ph3 was used to obtain the phase-contrast images with a 100x objective on an Olympus BX51 microscope.

**Synthesis of glycolipids**. All glycolipids described in this study were synthesized based on a small variation of previously published procedure (1). A solution of 1 equivalent of the unsaturated fatty acid and 1.1 equivalents of *N*, *N*-diisopropylethylamine (DIPEA) in dimethyl formamide (DMF) was stirred at 0 ºC for 10 min, and then 1.1 equivalents of **1-[Bis(dimethylamino)methylene]-1*H*-1,2,3-triazolo[4,5-*b*] pyridinium 3-oxide hexafluorophosphate (**HATU) (1.1 equivalents) was added. After 10 min stirring at 0 °C, 1 equivalent of β-D-glycopyranosylamine was added. After stirring for 1 h at room temperature, the solvents were removed *in vacuo* to obtain a yellow residue. This was washed with cold dilute HCl. The resulting off-white crude is dissolved in methanol and the slightly yellow solution is filtered using a 0.2 µm syringe-driven filter and purified by HPLC (Zorbax SB-C18 semipreparative column, 5% *Phase A* in *Phase B*, 6.5-7.5 min). The purified glycolipids are obtained as white powders.

**Droplet formation**. Typically, 9 µL of GOA from a 10 mM stock solution in methanol were added to 50 µL chloroform in a glass vial. When droplets were doped with amphiphiles bearing affinity handles, stock solutions were prepared in methanol and added into the chloroform as well. The organic solvents were evaporated under a slow stream of nitrogen gas and, while carefully rotating the glass vial, a thin film was generated at the bottom of the vial. The film was hydrated with a solution containing different amounts of IGEPAL (Sigma I8896, Lot # MKCC9036) within the droplet forming composition range (**Fig. 1*D***) as indicated for each experiment. The vial was vortex mixed until the lipid film was completely resuspended and the dispersion appeared turbid. To ensure complete hydration of the film, the sides and bottom of the vial are scratched with the pipette tip. As droplets form robustly at different GOA and IGEPAL concentrations and in different buffer compositions, the volume and composition of the rehydration solution was adjusted as indicated for each experiment.

**Thermogravimetric analysis**. A droplet dispersion was prepared in MilliQ $H_2O$ from 3 mM GOA and 2.25 mM IGEPAL and allowed to coalesce into a single phase. The coalesced phase (2.351 mg) was carefully collected from the bottom of the tube and loaded onto a ceramic crucible. Thermogravimetric analysis was carried out on a Perkin-Elmer STA 6000 Simultaneous Thermal Analyzer. The sample was heated at 10 °C/min over the range 30-350 °C. Nitrogen gas flow was maintained at 20 mL/min. We present the data over the range 30-150 °C. Above ~200 °C, the residual materials underwent thermal decomposition (data not shown).

**Cryogenic Transmission Electron Microscopy (Cryo-TEM)**. Sponge droplets were prepared from 3 mM GOA and 2.25 mM IGEPAL and sonicated with heating to ensure formation of small droplets. Immediately before grid preparation, the droplet sample was pipetted onto plasma-cleaned 300-mesh Quantifoil Multi-A carbon grids (Quantifoil). Using a Vitrobot EM grid plunger

(FEI), excess buffer was blotted at room temperature and 95% humidity and the grids were plunge-frozen in liquid ethane maintained at about -180 °C. The grids were stored in liquid nitrogen until use. Droplet samples were imaged on a JEM-2100F microscope (JOEL) fitted with a OneView 4K charge-coupled device camera (Gatan).

**Cryogenic Scanning Electron Microscopy (Cryo-SEM)**. Approximately 3 µL of sample (dispersion of 3 mM GOA and 2.25 mM IGEPAL) was pipetted onto a 3 mm Al planchette and flash frozen in liquid ethane before loading into a Leica cryo-transfer shuttle for insertion into a Leica ACE600 cooled down to -130 °C. The sample was then freeze-fractured and freeze-etched for 6 minutes at -105 °C and coated with 7 nm of C/Pt and an additional 5 nm of C. The sample was then transferred to in a Hitachi S-4800 cFEG-SEM fitted with a Leica cryo-stage cooled down to -130 °C. Imaging occurred at 5 kV with an approximate working distance of 9.7 mm.

**Small-angle X-ray scattering (SAXS)**. For studying the sponge phase characteristics, we prepared a series of samples with controlled ratios between GOA and IGEPAL. The droplet dispersions were loaded in thin walled (0.01 mm) special glass capillary tubes of 15 mm outer diameter (15-SG, Charles Supper Company). Synchrotron X-ray scattering measurements were performed at the Advanced Photon Source (APS), Argonne National Laboratory (ANL) at beamline 12-ID-C with monochromatic photon flux $1 \times 10^{13}$ photons/s/cm$^2$. A focused beam of dimension $1 \times 1$ mm$^2$ with incident photon energy at 18 keV (wavelength = 0.69 Å) was employed with exposure times of 10 s to get a reasonable scattering signal from the sample. The SAXS data were collected by a CCD detector (2048 × 2048 pixels) and the sample-to-detector distance was 3825 mm. To obtain one-dimensional SAXS profiles, 2-dimensional scattering data were azimuthally averaged with proper background subtraction. Finally, the data were transformed into the profiles of scattering intensity as a function of scattering vector ($q$).

**Sequestration experiments**. All droplet dispersions for the sequestration experiments were prepared from 3 mM GOA and 2.25 mM IGEPAL such that the total mass of droplet forming amphiphiles was 80 µg. 5 µg of the molecule to be sequestered were added in each case. In case of sequestration studies of fluorescein, POPC, and cholesterol, 5 µL of 1 mg/mL solution in EtOH were added during GOA film formation. 20 µL of the cloudy droplet dispersion was collected in a 0.6 mL Eppendorf tube and placed in an incubator at 37 $^{o}$C for 2 h. The tubes were centrifuged at 14,000 rcf for 2 min to ensure complete coalescence of the droplets. 2 µL of the supernatant was collected carefully and diluted with 98 µL 1:1 MeOH: $H_2O$. This was injected as whole into HPLC-MS and the peak area corresponding to the analyte in consideration was measured (denoted by $S$). The coalesced phase was resuspended by vigorous vortexing and 2 µL of the dispersion was diluted with 98 µL 1:1 MeOH: $H_2O$. This was injected as whole into HPLC-MS and the peak area corresponding to the analyte in consideration was measured (denoted by $T$). The sequestered fraction was calculated from the following equation:

$$Fraction\ Sequestered = 1 - S \div [S \times 0.1 + T \times 0.9]$$

The experiments were carried out in triplicate and the combined data are presented in ***SI Appendix*, Table S1**.

**Sequestration of LecA (PA-IL).** *Fluorescent labeling of LecA*. Terminal galactose-binding lectin LecA (or PA-IL) from *Pseudomonas aeruginosa* was purchased from Sigma-Aldrich. 0.31 mg of the solid powder (~80% protein) was dissolved in 200 µL 100 mM $NaHCO_3$, pH 8.3. 3.3 equivalents of Alexa Fluor 488 NHS ester (dissolved in anhydrous DMSO) was added to this solution, and the mixture was stirred at room temperature for 2 h. The excess dye was removed by a commercial dye removal kit (Pierce, Thermo Fisher Scientific). The eluent was further washed and exchanged with 1X PBS using a 10 kDa MWCO spin filter and stored at -20 $^{\circ}$C as 10% glycerol stock containing 0.02% (w/v) $NaN_3$.
*Microscopic observations*. GOA droplets were prepared with 3 mM GOA and 1.33-2.25 mM IGEPAL. Sequestration of various concentrations of Alexa Fluor 488 labeled LecA (1.0 µM-0.25 nM) was studied by spinning disk confocal microscopy.

**Reconstitution of Cytochrome c Oxidase (CcO)**. *Reduction of ferricytochrome c*. 1.72 mg of reddish-brown powder of ferricytochrome c from bovine heart (Sigma) were dissolved in 190 µL PBS (1X). Then, 10 µL of sodium ascorbate (200 mM) were added. The mixture was kept at 4 $_{o}$C overnight, observing that its color changed to rose-red. The residual ascorbate was removed by dialysis (mini dialysis device, 10 kDa MWCO, Slide-A-Lyzer$_{TM}$) against 10 mM sodium phosphate buffer (pH 7.0) followed by spin filtration using a 10 kDa MWCO spin filter with 6X volume. The final concentration of ferrocytochrome c was determined to be 1.11 mM and the $A_{550}/A_{565}$ ratio was 17.43.

*Detergent exchange of CcO.* 5 µL of CcO from bovine heart (~5 mg/mL in 39 mM DDM, 25 mM Tris pH 7.8, 5 mM EDTA) was added to 197.5 µL (in 10 mM sodium phosphate buffer, pH 7.0) of 1.68 mM IGEPAL. 2.5 µL of this solution was dissolved in 97.5 µL MeOH and mixed by vigorous vortexing. The tube was centrifuged at maximum speed to remove any insoluble material. The supernatant was carefully collected and run on HPLC-MS and the peaks corresponding to DDM and IGEPAL were seen distinctly. The remaining solution was transferred to a 100 kDa MWCO spin filter and thoroughly washed with (5×200 µL) of 1.68 mM IGEPAL (in 10 mM sodium phosphate buffer, pH 7.0). 1 µL of the final retentate (~35 µL) was added to 99 µL MeOH and run on HPLC-MS as described. The analysis verified that the DDM present in the commercially available CcO was completely exchanged with IGEPAL. Based on a calibration curve, the concentration of IGEPAL was estimated to be 18.8 mM.

*Activity assay of CcO*. A lipid film from 39 µL of 10 mM GOA was hydrated with a solution containing 377 µL sodium phosphate (10 mM, pH 7.0), 6 µL CcO (in 18.8 mM IGEPAL and 10 mM sodium phosphate pH 7.0), and 7 µL of IGEPAL (16.83 mM in 10 mM sodium phosphate pH 7.0). The turbid droplet dispersion was transferred to a cuvette and 15 µL of ferrocytochrome c was added. The dispersion was mixed by tapping and pipetting and placed in Nanodrop 2000c.

Absorbance at 550 nm was measured every 5 s over 30 min and a steady decrease was observed. To compare the activity of CcO at the same concentration of IGEPAL in the absence of droplets, a sample was prepared in an identical manner and the activity was assayed.

A negative control sample was prepared and assayed where no CcO was added. The $A_{550}$ was recorded identically and a negligible decrease was observed.

*Fluorescent labeling of CcO*. 60 µg of commercially available CcO from bovine heart (Sigma) was exchanged with sodium bicarbonate buffer (100 mM, pH 8.3) containing 0.4 mM *n*-dodecyl-*β*-maltoside (DDM). Then, the solution was treated with 10 equivalents of Alexa Fluor 488 NHS ester (dissolved in anhydrous DMSO), and the mixture was stirred at room temperature for 2 h. Then, the excess dye was removed and DDM was exchanged with IGEPAL by spin filtration (100 kDa MWCO).

**Activity of Cathepsin K in droplets**. *General*. Recombinant $His_6$-tagged Human Cathepsin K was purchased from EMD Millipore. The enzyme was stored as 18.5 µM aliquots in 100 mM NaOAc/HOAc buffer (pH 5.5) containing 1 mM TCEP at -80 °C. The protein was labeled with Alexa Fluor 488 and microscopy verified its sequestration into droplets doped with Ni-NTA-DGS. Benzyloxycarbonyl-L-leucyl-L-arginine 7-amido-4-methylcoumarin (Z-LR-AMC) was purchased from Peptides International and stored as 10.8 mM DMSO solution at -80 °C. It was estimated from HPLC that 75% of Z-LR-AMC gets partitioned to the droplets used in the subsequent experiments.

*Fluorometric assay*. Ni-NTA-DGS/GOA/IGEPAL (26.7 µM/0.96 mM/0.64 mM) droplets were prepared in 100 mM HEPES-Na containing 1.5 mM TCEP. $His_6$-Cathepsin K was added to the droplet dispersion so that the enzyme concentration with respect to bulk was 2.3 nM. After incubating at room temperature for 10 min, Z-LR-AMC (dissolved in DMSO) was added such that the final concentration was 81 µM. Additional conditions were tested where one or more of the components were omitted: GOA/IGEPAL (0.96 mM/0.64 mM), IGEPAL (0.64 mM), bulk, and negative control (no amphiphiles and no enzyme). 20 µL of each sample was loaded onto a 384 well-plate and analyzed for fluorescence from 7-amino-4-methylcoumarin ($\lambda_{ex}$: 360 nm, $\lambda_{em}$: 460 nm; 10 nm bandwidth) every 1 min at room temperature in a Tecan Spark microplate reader.

*HPLC-MS assay*. Calibration curves were prepared for Z-LR-AMC and AMC with known quantities of the pure substances (***SI Appendix,*** **Fig. S14**). Ni-NTA-DGS/GOA/IGEPAL (26.7 µM/0.96 mM/0.64 mM) droplets were prepared in 100 mM HEPES-Na containing 2.0 mM TCEP. $His_6$-Cathepsin K was added to the droplet dispersion to a bulk concentration of 2.8 nM. Z-LR-AMC was added to a final concentration of 81 µM. GOA/IGEPAL (0.96 mM/0.64 mM), IGEPAL (0.64 mM), bulk, and negative control (no amphiphiles and no enzyme) samples were also prepared in identical buffer and same quantity of enzyme (except for the negative control) and substrate were used. The reactions (20 µL) were incubated at room temperature for 6 h and quenched by addition of 80 µL of 1:1 MeOH:$H_2O$. The reactions were analyzed by HPLC-MS and the areas of the peaks under 330 nm chromatogram were used for calculation of yields. The experiments were repeated in triplicate.

**FRAP experiments**. For the analysis of lipid diffusion, droplets were formed with 3.6 mM GOA, 2.56 mM IGEPAL and 36 µM Texas Red DHPE in 1X PBS. For the analysis of protein diffusion, droplets were formed with 3 mM GOA, 2.25 mM IGEPAL and 83 µM Ni-NTA-DGS in 100 mM HEPES pH 8. Purified mCherry-$His_6$ protein (Antibodies-online Inc.) was added to a final concentration of 10 µM. As controls, we performed FRAP experiments on mCherry-$His_6$ bound to Ni-NTA agarose resin (HisPur, Thermo Fisher Scientific) and a water-in-oil emulsion of 90 µM mCherry-$His_6$ in mineral oil containing 0.25% (*v*/*v*) Span-80. FRAP experiments were performed with a Zeiss Spinning Disk microscope using a 63x 1.4 NA oil objective. A circular region of interest (ROI) with a diameter of 9.6 µm was bleached in large droplets (>50 µm) with a 5 s pulse of the 561 nm laser at 100% intensity. Samples were imaged every 500 ms with a 150 ms exposure at 50% intensity. Movies were analyzed in Fiji/ImageJ (2) to extract average fluorescence over time of the bleached ROI, entire droplet and background. Background corrected fluorescence values were used to correct for possible photobleaching

$$f(t) = \frac{F_{ROI}(t)}{F_{Whole}(t)}$$

and then scaled between 0 and 1

$$F(t) = \frac{f(t) - f(0)}{f(-10s) - f(0)}$$

where f(0) is the intensity at time 0s, right after the bleach, and f(-10s) is the initial prebleach intensity (3). Data were then fitted to

$$F(t) = A(1 - e^{-\frac{t}{\tau}})$$

where A is the amplitude of recovery and $\tau$ is the time constant that allowed us to calculate the half-time to recovery $t_{1/2}$.

$$t_{1/2} = ln2 \cdot \tau$$

**TX-TL in the presence of lipid sponge droplets**. A lipid film was prepared from 18 µL 10 mM GOA and rehydrated by vortexing with 9 µL rehydration solution (6 µL PURExpress Solution A, 0.375 µL Murine RNase Inhibitor (NEB), 1.5 µL 67.3 mM IGEPAL, 1.125 µL $H_2O$). For the reaction to start, 4.5 µL PURExpress Solution B and linear DNA templates were added for a final volume of 15 µL. DNA template concentrations in the TX-TL reaction were 5 nM for $P_{T7}$-sfGFP-lecA and $P_{T7}$-sfGFP, and 10 nM for $P_{T7}$-sfGFP-lacI. Linear DNA templates were assembled by PCR and PCR purified, their sequences are listed in ***SI Appendix,*** **Table S2**. Expression reactions were carried out at 33 °C, in droplets of 5 µL volume sandwiched between cover glass and gas permeable Lumox plates (Sarstedt) or in chambers of a microfluidic device (4), where fluorescence increase of individual droplets could be followed by time lapse imaging. For expression of DAGK, a lipid film was prepared from 15 µL of 10 mM GOA and 3 µL of 10 mM 1-decanoyl-*rac*-glycerol (Cayman Chemicals). Hydration was carried out with 30 µL of a solution containing 5 mM IGEPAL, 0.5 mM ATP, and 0.5 mM $MgCl_2$. Protein expression reaction was consisted of 8 µL PURExpress Solution A, 6 µL PURExpress Solution B, 0.5 µL Murine RNase Inhibitor (NEB), 1 µL DAGK linear DNA (28.8 nM final concentration), 1 µL $H_2O$, and 4 µL droplet dispersion (containing the substrates for DAGK). 10.25 µL of the reaction is taken out immediately and quenched with 4 µL $Na_2$-EDTA (100 mM) and 90 µL methanol. The remainder is incubated for 3 h at 37 °C and quenched similarly. The initial and final time points were analyzed by HPLC-MS in selective ion mode for *m*/*z* 269 (positive ion), and *m*/*z* 325 (negative ion). For the expression of sfGFP-DAGK, 13.4 nM of DNA template was used.

**Protein expression, purification and analysis**. Proteins (***SI Appendix,*** **Table S2,** ***SI Appendix,*** **Fig. S21**) were expressed in *E. coli* BL21 (DE3) using the plasmids listed in ***SI Appendix,*** **Table S3**. ClpX-$His_6$ and ClpP-$His_6$ were gifts from Xingying Shi and Simpson Joseph (UC San Diego) (5). Biotinylated PhyB(1-651)-AviTag with phycocyanobilin was produced in *E. coli* (6). Proteins were purified by Ni-NTA affinity chromatography. Purified proteins were buffer exchanged to storage buffer using spin filter devices or dialysis. Storage conditions were 200 mM potassium phosphate pH 7.2, 2 mM EDTA, 400 mM NaCl, 1 mM DTT, 1 mM $NaN_3$, 50% (v/v) glycerol at -20 °C for sfGFP-LacI-$His_6$, 1X PBS at -80 °C for sfGFP-$His_6$, 100 mM HEPES pH 8.0, 25 mM NaCl, 0.5 mM TCEP, 50% (v/v) glycerol at -20 °C for PhyB-AviTag, 100 mM HEPES pH 8.0, 25 mM NaCl, 5 mM DTT, 50% (v/v) glycerol at -20 °C for $His_6$-PIF-sfGFP-ssrA, and 50 mM Tris pH 7.5, 200 mM KCl, 25 mM $MgCl_2$, 0.67 mM IGEPAL, 0.5 mM TCEP, 20% (v/v) glycerol at -80 °C for ClpP-msfGFP-$His_6$. Protein concentrations were measured by BCA assay. PhyB-AviTag was verified by Zinc staining to visualize the bound chromophore phycocyanobilin by incubating an SDS-PAGE gel in 1 mM zinc acetate for 15 min followed by visualization under UV light. Photoresponsive switching was verified by recording absorbance spectra of purified PhyB protein after exposure to 660 nm or 740 nm light using a Tecan Spark microplate reader. For immunoblotting, protein samples were separated by SDS-PAGE and transferred to PVDF membranes. Antibodies used were GFP Antibody (B-2) (Santa Cruz Biotechnology) at 1:200 and 6x-His Tag Monoclonal Antibody (Invitrogen) at 1:1000. Bands were detected with a horseradish peroxidase coupled secondary antibody and SuperSignal West Pico PLUS Chemiluminescent Substrate (Thermo Fisher Scientific).

**Light-control of protein localization**. Samples were illuminated using LEDs (Roithner LaserTechnik GmbH, LED660N-03 and LED740-series) in light insulated containers, where LEDs were placed in proximity to samples using a custom-built setup. In 740 nm light conditions, samples were continuously illuminated by 740 nm light. In 660 nm light conditions, samples were illuminated every 10 min with a 30 s light pulse and otherwise incubated in the dark.

**ClpXP reactions**. For light controlled protein degradation, droplets for one ClpXP reaction were prepared as follows, where the volumes were scaled according to the reactions performed in one experiment. Per reaction, a lipid film was formed from 7.2 µL 10 mM GOA and 0.8 µL 2.5 mM 18:1 Biotinyl-PE (Avanti). The lipid film was vortexed in 40 µL rehydration solution (1.35 mM IGEPAL in 100 mM HEPES pH 8.0) to form droplets. 40 µL of 2 µM Streptavidin and 2 µM PhyB-AviTag in reaction buffer [50 mM HEPES pH 8.0, 20 mM KCl, 5 mM $MgCl_2$, 1% (v/v) glycerol, 1% (w/v) BSA, 5 mM β-ME, 0.067 mM IGEPAL] were added and incubated with droplets for 15 min. Unbound PhyB and Streptavidin were then removed by washing with reaction buffer by pelleting droplets by centrifugation at 1500 rcf. During washing, droplets were concentrated in 10 µL reaction buffer, and the substrate PIF-sfGFP-ssrA was added for a final concentration of 250 nM in the reaction for binding to PhyB-loaded droplets and incubated for 15 min. The reaction was started by adding premixed ClpXP and ATP in reaction buffer (concentrations were 4 mM ATP, 50 nM $ClpX_6$, 200 nM $ClpP_{14}$ in a final reaction volume of 20 µL), and droplets were coalesced in the bottom of the reaction plate (384-well v-bottom plates, Promega) by centrifugation. The reaction was performed at room temperature, monitored by reading fluorescence of PIF-sfGFP-ssrA in a Tecan Spark microplate reader (excitation 485 +/- 20 nm (filter), emission 520 nm +/- 20 (monochromator)) every ten minutes and placed in specific illumination conditions between reads. For analysis by western blot, samples were collected at different time points, quenched by adding Laemmli buffer and stored at -20 °C. For analysis by microscopy, samples were first placed in 740 nm from 5 min and then in 660 nm light for 30 min to compare droplet fluorescence.

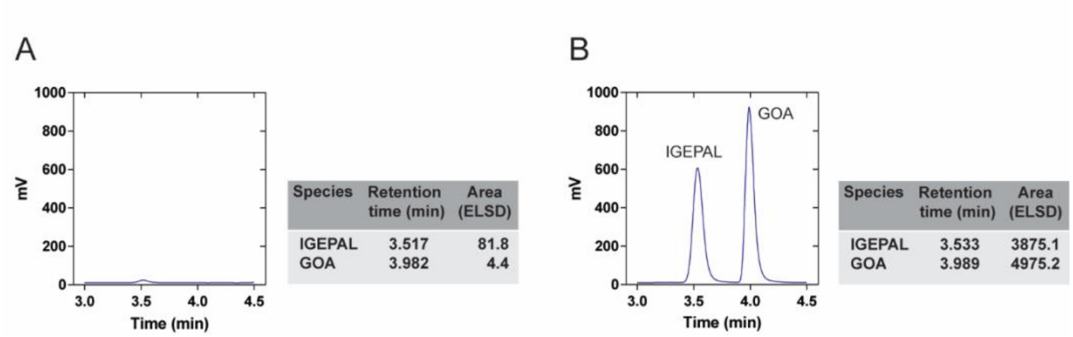


**Fig. S1.** Estimation of the concentration of amphiphiles in the sponge phase by HPLC. A representative ELSD chromatogram comparing the compositions of (*A*) supernatant and (*B*) resuspended coalesced phase from a droplet sample composed of 2.13:1 GOA:IGEPAL (by molar ratio).

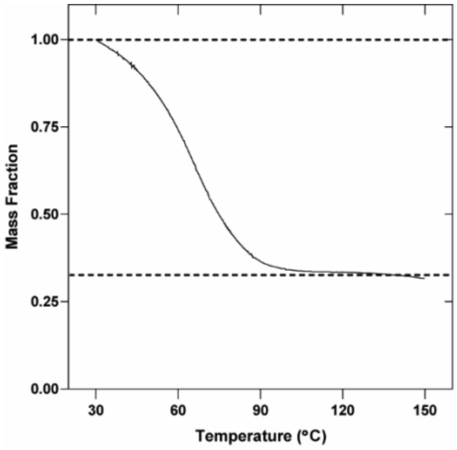


**Fig. S2.** Estimation of the water content in the sponge phase by Thermogravimetric Analysis (TGA). TGA trace obtained when a GOA:IGEPAL (4:3 molar ratio) coalesced phase was heated. The dotted lines are meant to provide a guide to the eye.

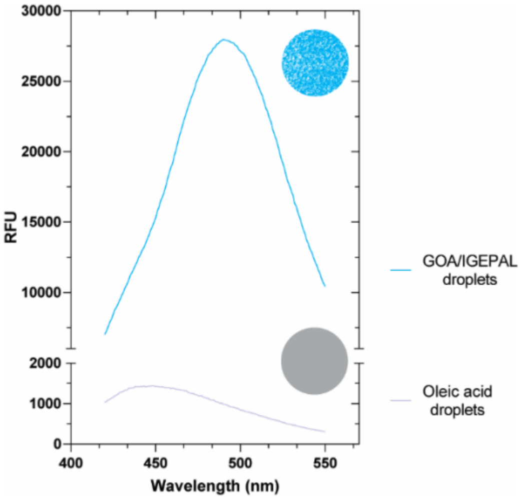


**Fig. S3.** Fluorescence properties of Laurdan in sponge phase. Fluorescence spectrum of the solvatochromic dye Laurdan (0.3 mol%) in sponge phase shows a strong maximum at 490 nm suggesting the presence of a water-accessible fluid environment inside the droplets (7). In comparison, Laurdan shows a maximum at 440 nm in oleic acid oil droplets.

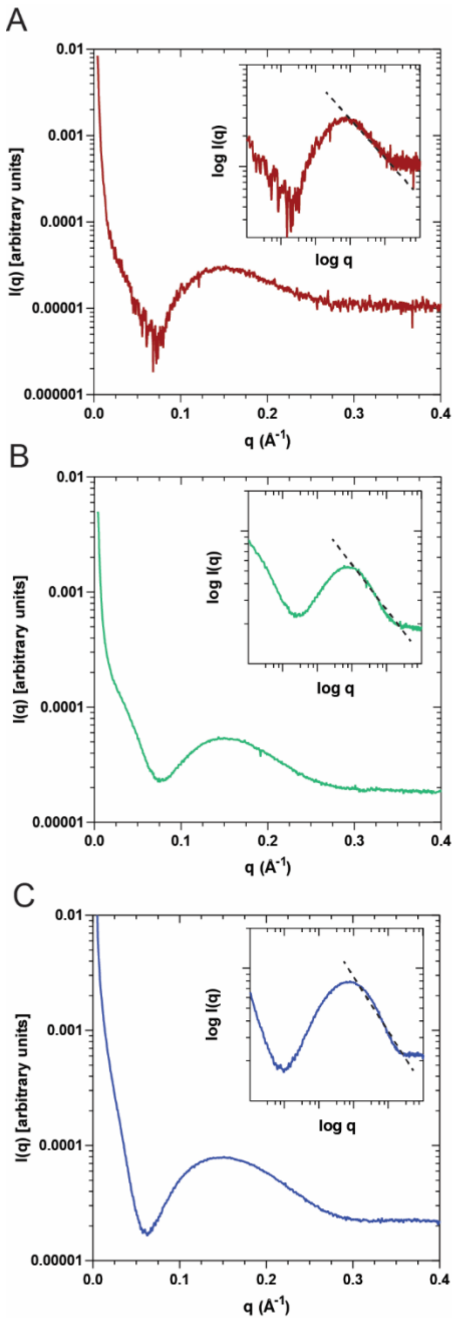


**Fig. S4.** SAXS intensity profiles of sponge phase show characteristics of a bilayer structure. SAXS intensity profiles from GOA:IGEPAL droplet dispersions in the molar ratio (*A*) 1:1 (*B*) 3:2 (*C*). 3:1 shows a $I(q) \approx q_{-2}$ decay, suggesting a local bilayer structure (8–10). The corresponding log-log plot is shown in the inset. The slopes are calculated to be 1.90, 1.96, and 2.18 respectively.

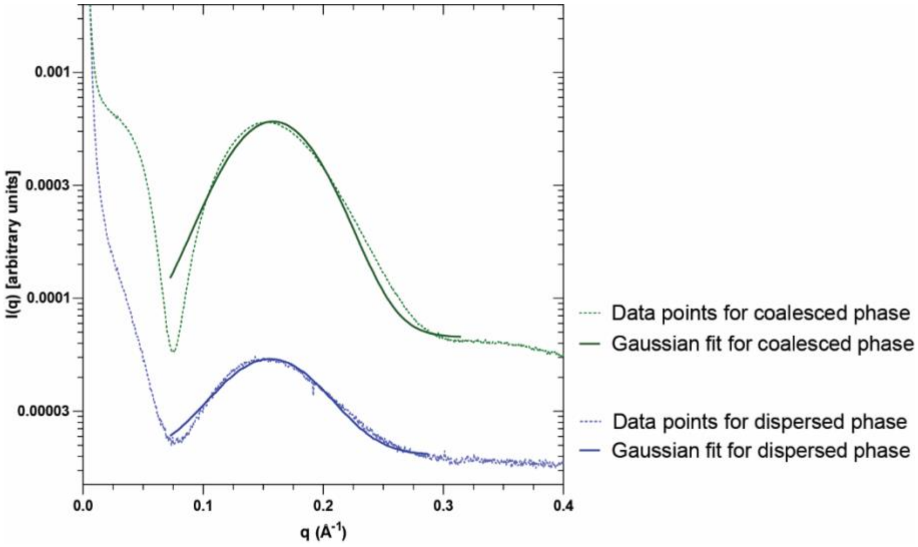


**Fig. S5.** SAXS intensity profiles of dispersed and coalesced phases. When the SAXS intensity profiles of dispersed and coalesced phases from 3:2 GOA:IGEPAL (by molar ratio) are compared, the nature of the profiles and position of maxima are observed to be practically unchanged. The solid lines correspond to the Gaussian distribution fit to the sponge phase peak data points. For dispersed phase, $q_{max}$ = 0.153±0.0002 Å$^{-1}$, for coalesced phase $q_{max}$ = 0.156±0.0002 Å$^{-1}$.

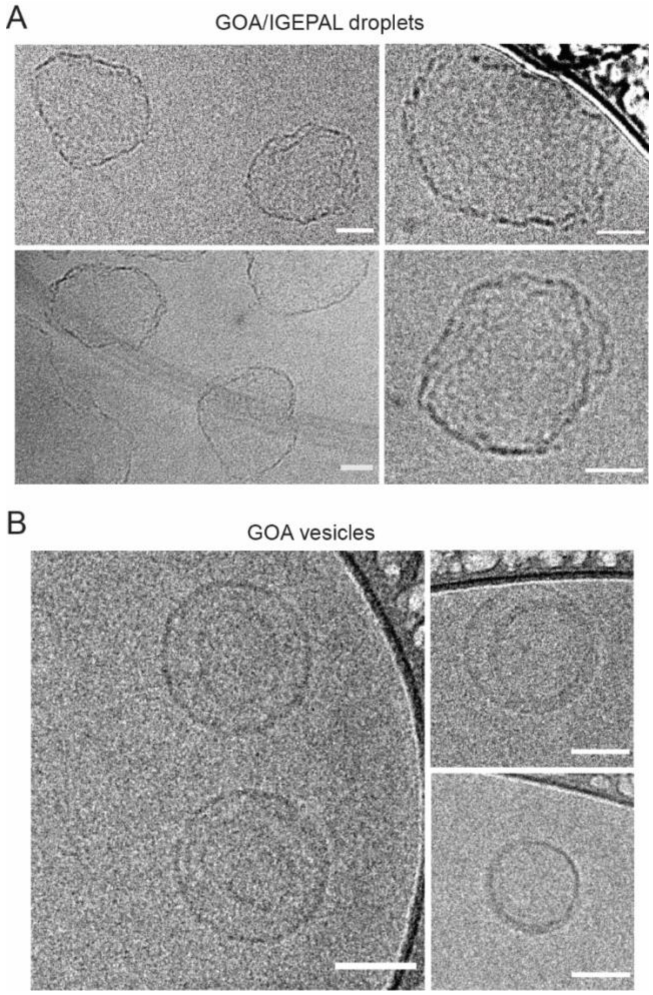


**Fig. S6**. Cryogenic transmission electron microscopy on lipid sponge droplets and vesicles. (*A*) Sonicated sponge droplets from GOA and IGEPAL. (*B*) Sonicated GOA vesicles. All scale bars: 50 nm.

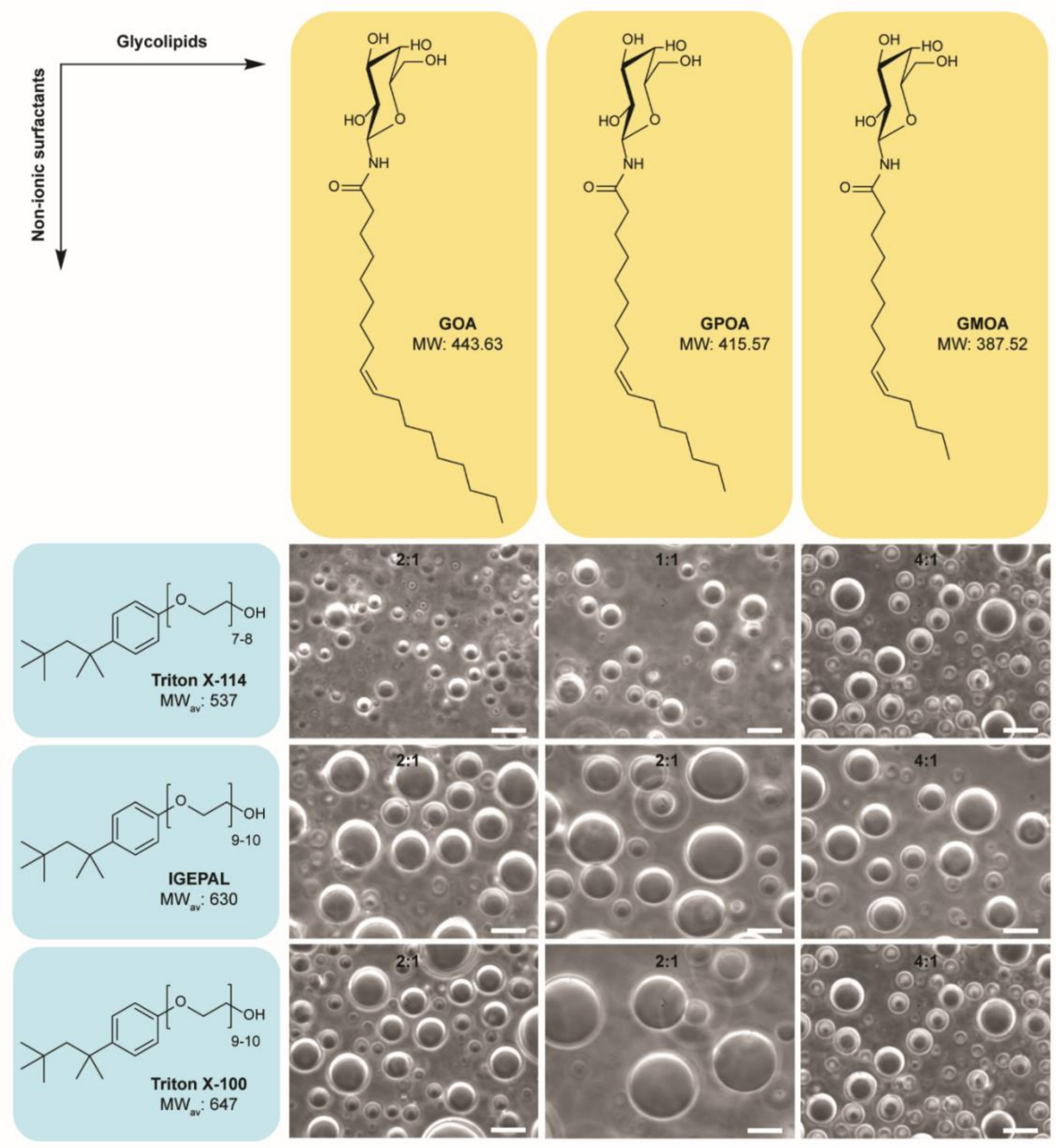


**Fig. S7**. Droplet formation from various combinations of glycolipids and non-ionic detergents. Phase contrast microscopy of droplets formed from various combinations of β-D-galactopyranosylamides of unsaturated fatty acids (1) and octylphenoxypolyethoxyethanol surfactants in 1X PBS. The ratios on the images correspond to those of the glycolipid and the detergent used. All scale bars: 10 µm.

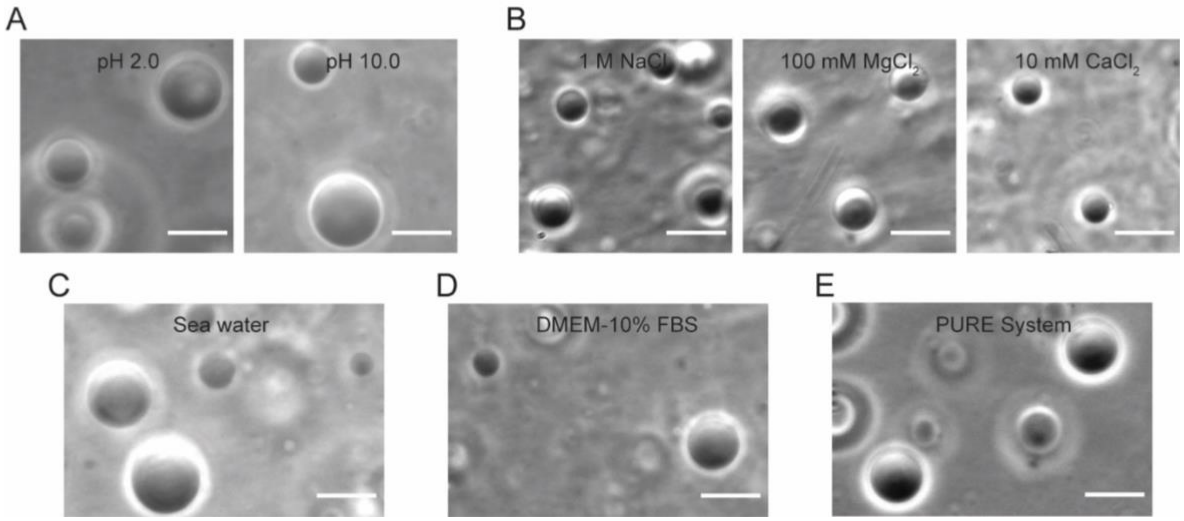


**Fig. S8.** Robustness of droplet formation. Formation of droplets from GOA and IGEPAL at (*A*) pH 2.0 (Na-glycine buffer), pH 10.0 (Na-glycine buffer) (*B*) 1 M NaCl, 100 mM $MgCl_2$, 10 mM $CaCl_2$ (*C*) sea water (*D*) DMEM-10% fetal bovine serum (FBS) (*E*) PURE system (cell free transcription-translation medium). All scale bars: 10 µm.

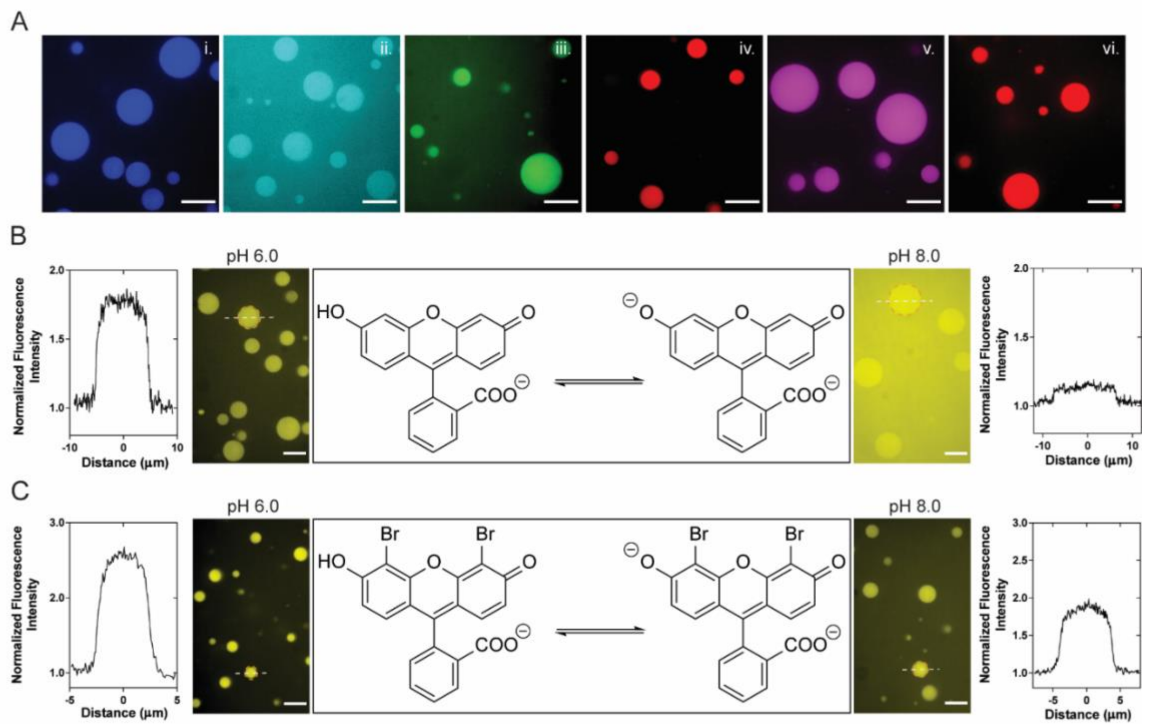


**Fig. S9.** Partitioning of fluorescent molecules into the sponge phase. (*A*) Confocal microscopy of sponge droplets partitioning i. 7-amino-4-methylcoumarin ii. 7-hydroxycoumarin iii. Rhodamine Green iv. Rhodamine X v. Rhodamine 6G and vi. Sulforhodamine B. All scale bars: 15 µm. Extent of partitioning of (*B*) fluorescein and (*C*) 4´, 5´-dibromofluorescein into sponge droplets changes with pH (6.0, 8.0). All scale bars: 10 µm. The adjoining intensity profile plots correspond to the circled droplets. On *x*-axes, ‘0’ corresponds to the center of the droplet.

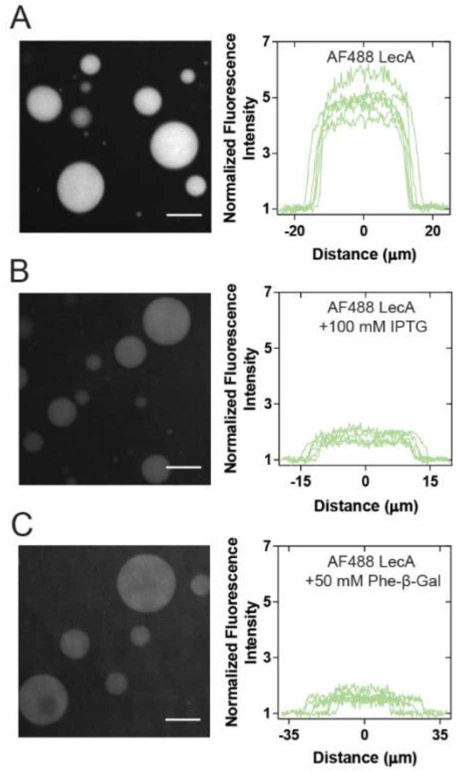


**Fig. S10.** Partitioning of Alexa Fluor 488-labeled LecA in presence of non-amphiphilic competitive ligands. Droplets are prepared from 3 mM GOA and 1.8 mM IGEPAL in 1X PBS and 1.5 µM Alexa Fluor 488-labeled LecA is added in presence of (*A*) no competitive ligands (*B*) 100 mM IPTG (*C*) 50 mM phenyl β-D-galactopyranoside. All fluorescence images are window-leveled identically. All scale bars: 20 µm. The adjacent plots correspond to the fluorescence intensity across the middle of 6 droplets. The ‘0’ on *x*-axes corresponds to the center of the droplet.

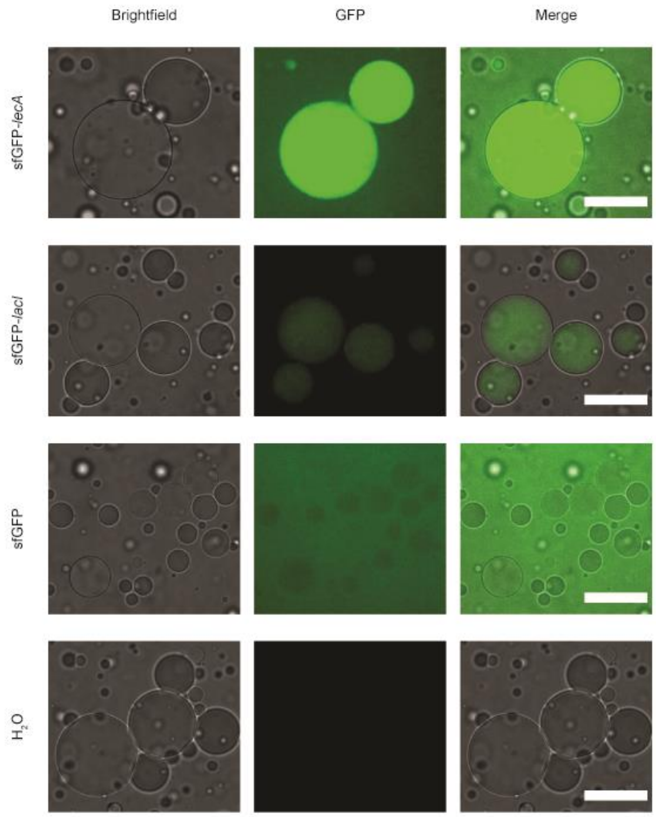


**Fig. S11.** In vitro transcription and translation in the presence of lipid sponge droplets. Proteins were expressed in a recombinant cell-free expression system (PURExpress, New England Biolabs) from linear DNA under control of the T7 promoter in the presence of droplets. Images were acquired after 3 h and are displayed with the same contrast settings. Scale bars: 50 µm.

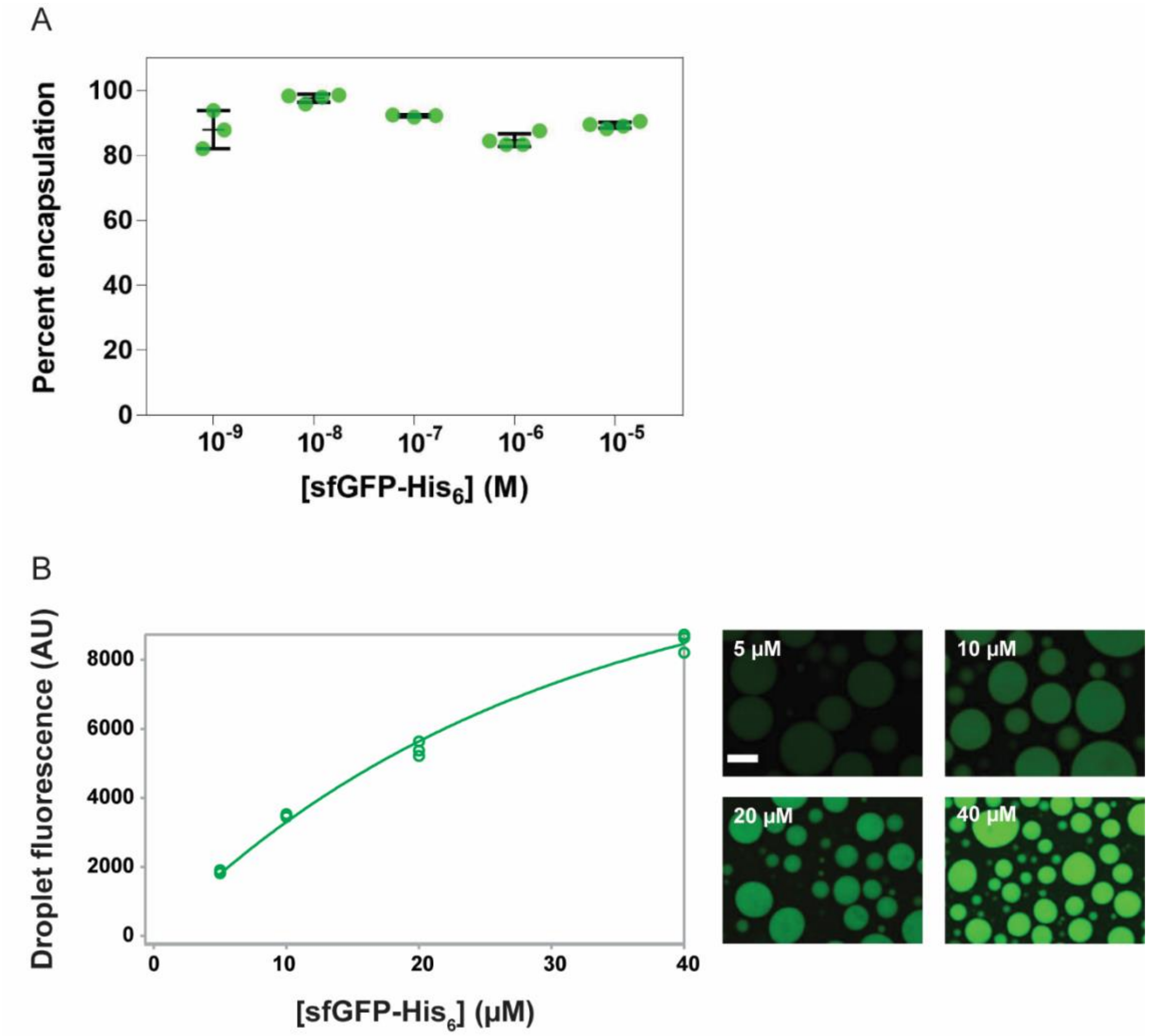


**Fig. S12.** Partitioning of sfGFP-$His_6$ in Ni-NTA-DGS doped sponge droplets. (*A*) Quantification of the extent of partitioning of sfGFP-$His_6$ into sponge phase droplets doped with 1.6 mol% Ni-NTA-DGS at various concentrations of the protein. Partitioning was estimated from fluorescence of the supernatant. (*B*) Partitioning measured by confocal microscopy. Droplet fluorescence intensities at increasing sfGFP-$His_6$ concentrations were measured in three images and data fitted to an exponential saturation function. Fluorescence begins to saturate as protein concentration approaches total Ni-NTA-DGS concentration (83.3 μM). Shown are representative images of droplet fluorescence displayed at identical contrast settings. Scale bar: 25 μm.

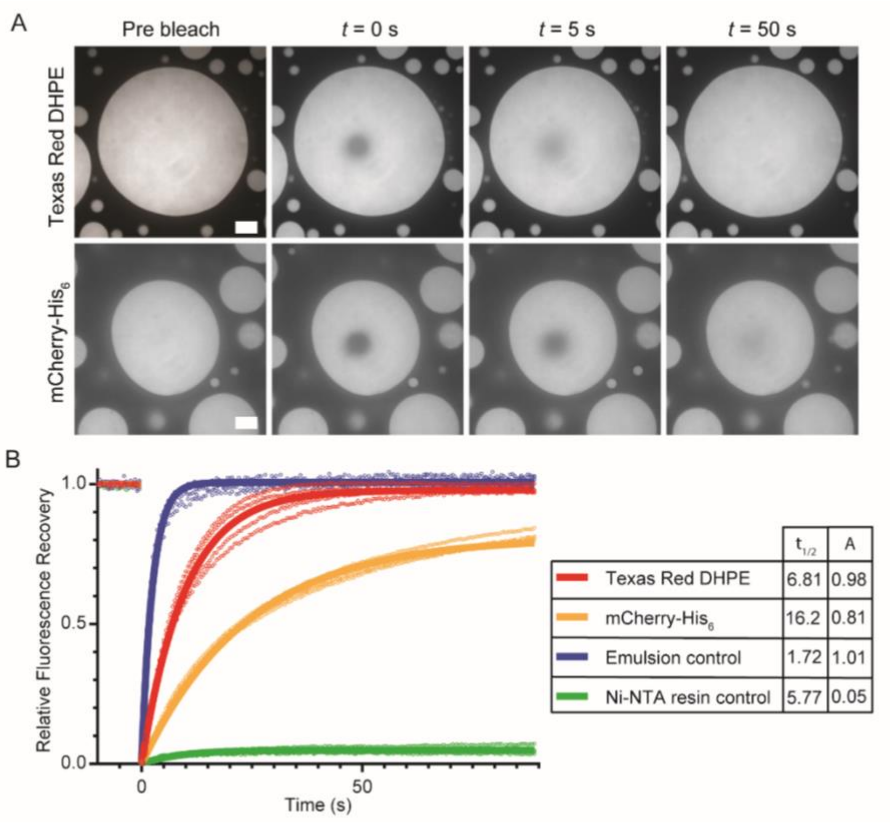


**Fig. S13.** FRAP analysis to characterize diffusion of biomolecules in lipid sponge droplets. (*A*) Representative fluorescence images of FRAP experiments on lipid sponge droplets containing the fluorescent phospholipid Texas Red-DHPE or fluorescent protein mCherry-$His_6$. Scale bars: 10 µm. (*B*) Relative fluorescence recovery of Texas Red DHPE and mCherry-$His_6$ in droplets, compared to fluorescence recovery of mCherry-$His_6$ in water-in-oil emulsion droplets and bound to a solid agarose Ni-NTA resin. Shown are data points from at least three FRAP experiments. The fraction of recovery (plateau), A, and $t_{1/2}$ (time to half-maximal recovery, in s) were obtained by fitting the data (solid lines) as described in the Materials and Methods section.

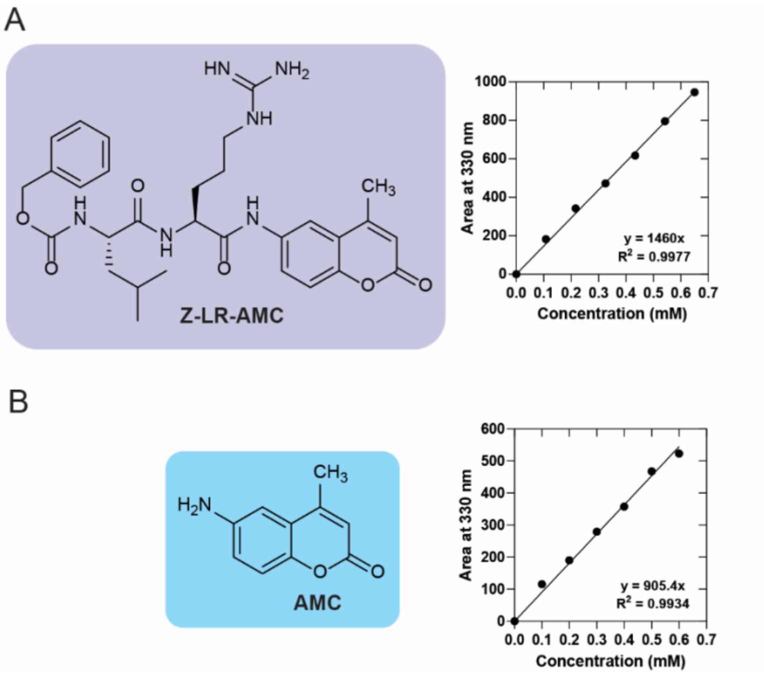


**Fig. S14.** HPLC calibration curves for Cathepsin K experiments. (*A*) benzyloxycarbonyl-L-leucyl-L-arginine 7-amido-4-methylcoumarin (Z-LR-AMC) (*B*) 7-amino-4-methylcoumarin (AMC). The areas under the 330 nm chromatogram were integrated for known amounts of both analytes to generate the calibration curves.

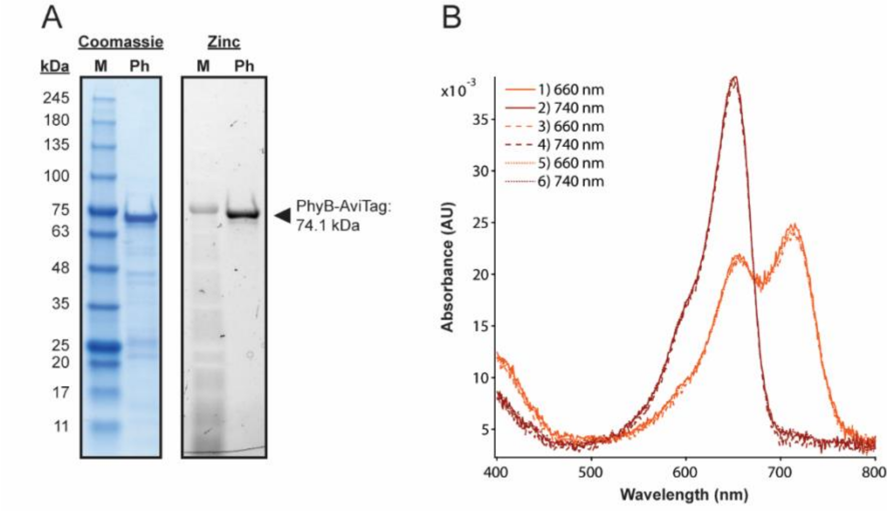


**Fig. S15.** Characterization of PhyB. (*A*) Coomassie and Zinc staining of purified PhyB-AviTag (2 µg) after SDS-PAGE. Zinc staining visualizes the bound chromophore phycocyanobilin. (*B*) Absorbance spectra of purified PhyB-AviTag after alternating exposures to 30 s of 660 nm and 740 nm light.

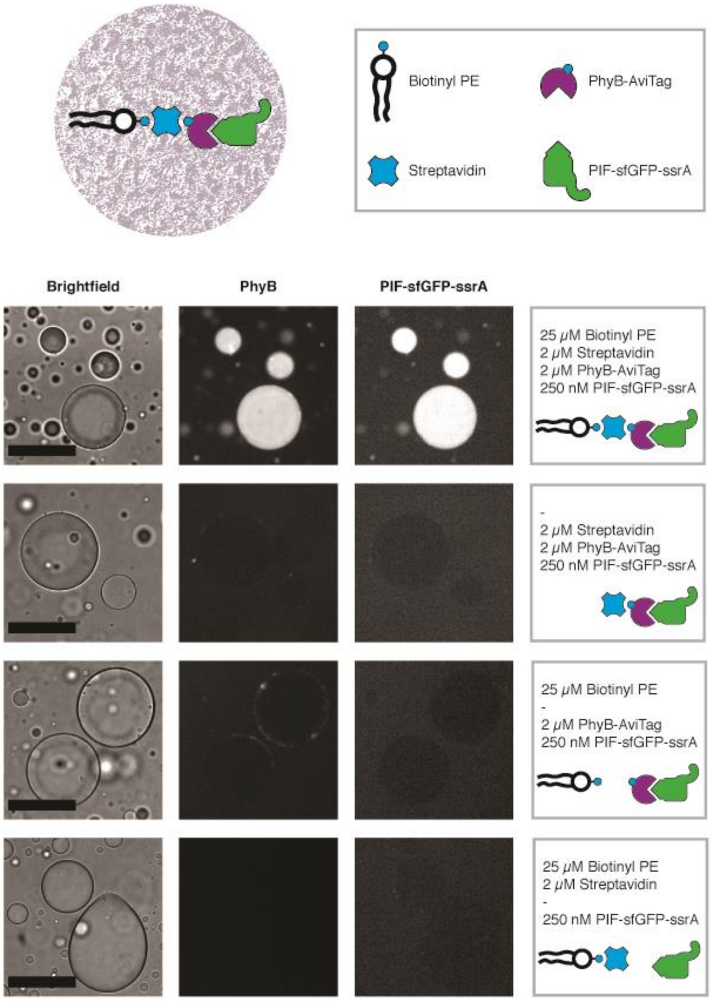


**Fig. S16.** Encapsulation of PhyB-AviTag and PIF-sfGFP-ssrA in lipid sponge droplets. Biotinyl PE and Streptavidin mediated encapsulation of biotinylated PhyB-AviTag and PIF-sfGFP-ssrA. For controls, individual components of the system were omitted as indicated on the right. PhyB-AviTag was visualized in the Cy5-channel. All images are displayed with the same contrast settings. Scale bars: 50 µm.

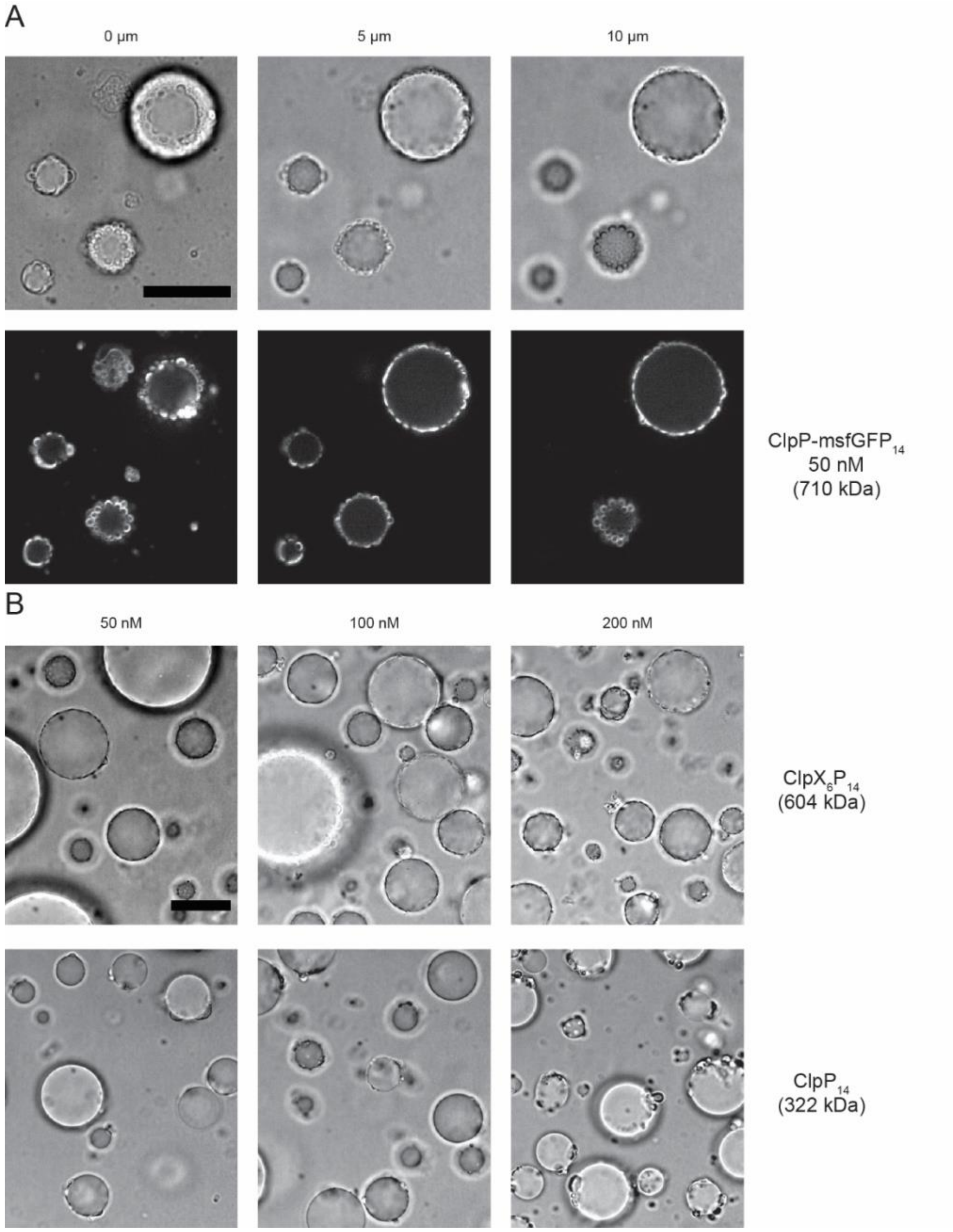


**Fig. S17**. Exclusion of $His_6$-tagged ClpXP from Ni-NTA-DGS-doped droplets. (*A*) ClpP-msfGFP-$His_6$ aggregates around Ni-NTA-DGS-doped droplets forming a membranous corona around droplets. 50 nM ClpP-msfGFP-$His_6$ (14-mer concentration) was added to droplets and incubated for 2 h. Shown are images from different slices of a *z*-stack to visualize membrane formation at three different heights. (*B*)Titrating increasing concentrations of unlabeled $ClpX_6P_{14}$ and $ClpP_{14}$ leads to increasing aggregation defects on droplets. All ClpP and ClpX monomers were tagged with C-terminal $His_6$-tags. Similar defects were not observed for sfGFP-$His_6$ (28 kDa) that partitioned into Ni-NTA-DGS-doped droplets (Fig. 2*E*). Droplets were imaged after 2 h incubation in the presence of protein. Scale bars: 20 µm.

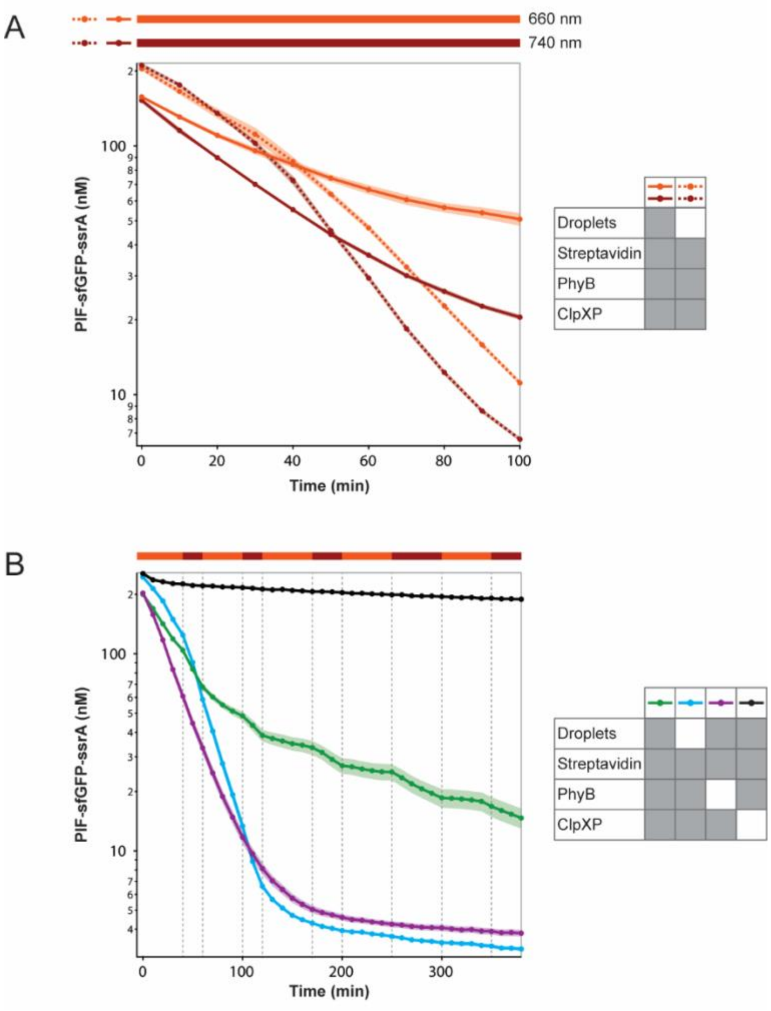


**Fig. S18.** Light-controlled protein degradation measured by fluorescence decrease. (*A*) Incubation of ClpXP reaction in 660 nm or 740 nm illumination conditions. In control reactions, droplets were omitted to analyze the effects of substrate binding to the PhyB-AviTag – Streptavidin complex alone. (*B*) Comparison to control reactions where individual reaction components were omitted. Switching of degradation rate in response to wavelength only occurs in the presence of all reaction components. All samples were illuminated by 660 nm or 740 nm light as indicated in the bar above the graph. Reactions were performed in duplicates (shaded area) with averages shown by solid markers.

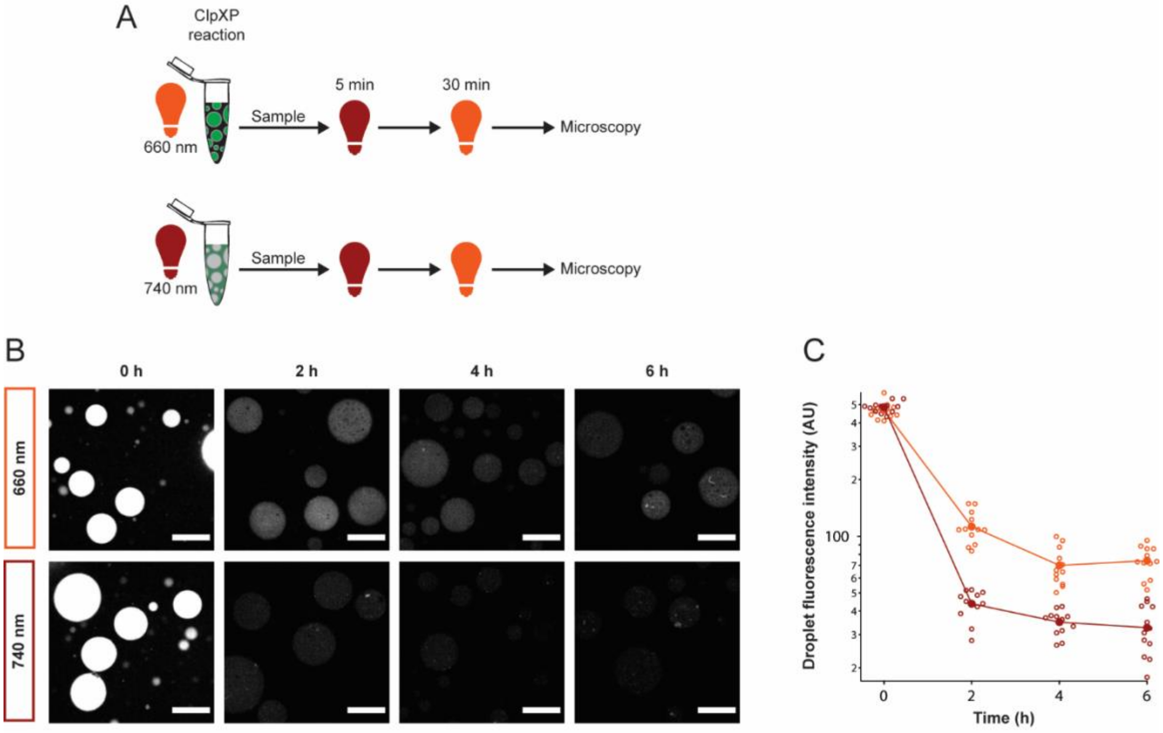


**Fig. S19**. Microscopy of light-controlled protection of PIF-sfGFP-ssrA in lipid sponge droplets. (*A*) Schematic of sample preparation. PhyB-loaded droplets were incubated with PIF-sfGFP-ssrA and ClpXP in 660 nm or 740 nm light. At different time points, samples were first placed in 740 nm for 5 min and then in 660 nm light for 30 min to compare droplet fluorescence by microscopy. (*B*) Representative images are displayed with the same contrast settings. Scale bars: 50 µm. (*C*) Quantification of droplet fluorescence from images. Droplet fluorescence of individual droplets was measured (and corrected for background fluorescence) in at least three images per condition and time point. Lines and solid marker: average.

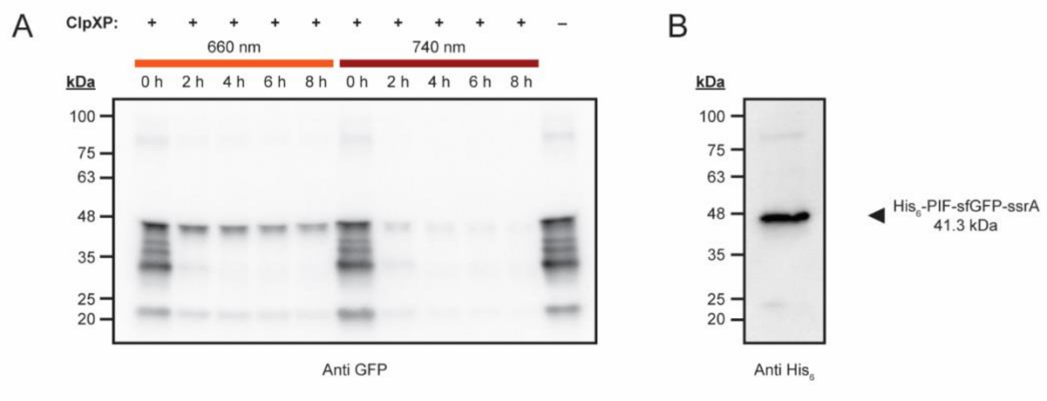


**Fig. S20.** Immunoblot analysis of light-controlled protection of PIF-sfGFP-ssrA in lipid sponge droplets. (*A*) PhyB-loaded droplets were incubated with PIF-sfGFP-ssrA and ClpXP in 660 nm or 740 nm light for the indicated time lengths. A control sample (left lane) did not contain ClpXP protease. The 41.3 kDa full length $His_6$-PIF-sfGFP-ssrA fusion protein is clearly protected from degradation by binding in glycolipid sponge droplets in 660 nm light. The primary antibody against GFP reveals truncated fragments of the fusion protein that are degraded at comparable rates in the two wavelengths of light. (*B*) Immunoblot of pure PIF-sfGFP-ssrA with a primary antibody against the N-terminal $His_6$ tag shows that the protein fragments between 25 and 40 kDa in (*A*) are likely C-terminal fragments that do not contain the N-terminal $His_6$ and the PhyB-binding PIF-tag. These protein fragments can therefore serve as internal controls for temperature or other potential effects.

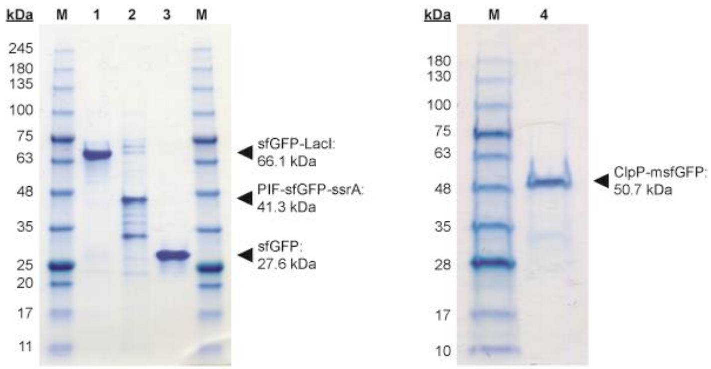


**Fig. S21.** SDS-PAGE analysis of purified sfGFP fusion proteins. Coomassie staining of Ni-NTA-purified proteins after SDS-PAGE (2 µg per lane were loaded on left gel, 1 µg per lane for ClpP-msfGFP). Lanes: **1** – sfGFP-LacI, **2** – PIF-sfGFP-ssrA, **3** – sfGFP, **4** – ClpP-msfGFP. **M**: Protein ladder.

**Table S1**. Summary of extent of partitioning of molecules (<1 kDa) into sponge phase. In each of these experiments, 5 µg of a molecule of interest was tested for partitioning in 80 µg of droplet material (composed of 4:3 GOA:IGEPAL by molar ratio) in 1X PBS. Using HPLC, the concentration of the molecule of interest in the supernatant was measured and then compared with the total to estimate the extent of partitioning. We define "capacity" as the mass of a molecule of interested partitioned per unit mass of droplet-forming amphiphiles.

**Partitioning of various molecules into lipid sponge phase**

| Molecule | Molar mass (g mol$^{-1}$) | Charge | Percentage partitioned | Capacity (mg g$^{-1}$) |
|---|---|---|---|---|
| Rhodamine B | 479.02 | 0 | 81.4±5.2 | 50.9 |
| Rhodamine 6G | 479.02 | +2 | 70.2±2.3 | 43.9 |
| Sulforhodamine B | 558.67 | -1 | 47.3±1.9 | 29.6 |
| 7-Diethylamino-4-methylcoumarin | 231.29 | +1 | 80.6±1.3 | 50.4 |
| 7-Amino-4-methylcoumarin | 175.18 | 0 | 73.7±5.5 | 46.1 |
| 7-Hydroxycoumarin (pH 7.4) | 162.14 | -1 | 6.3±0.9 | 3.9 |
| Fluorescein (pH 6.0) | 332.21 | -1 | 43.7±1.2 | 27.3 |
| Fluorescein (pH 8.0) | | -2 | 13.2±3.4 | 8.3 |
| Doxorubicin (pH 7.4) | 543.52 | +1 | 24.7±7.2 | 15.4 |
| Cholesterol | 386.65 | 0 | 100±0 | 62.5 |
| POPC | 760.09 | 0 | 100±0 | 62.5 |
| Adenosine | 267.24 | 0 | <1 | -- |
| 5´-*N*-Ethylcarboxamidoadenosine | 308.29 | 0 | <1 | -- |

**Table S2.** Sequences of linear DNA templates for TX-TL used in this study

| **Linear template** | **Annotation of features and DNA sequence** |
|---|---|
| $P_{T7}$-sfGFP-*lecA* | T7 promoter, ribosomal binding site, sfGFP, *lecA*, T7 terminator |
| | GTCTTCACCTCGAGGATCTTAAGGCTAGAGTAATACGACTCACTATAGGGAGA<br>TGTGGTCTAGACATTCCAGGTTAAGAAGGAGGAAAAAAAAATGCGTAAAGGA<br>GAAGAACTTTTCACTGGTGTCGTCCCTATTCTGGTGGAACTGGATGGTGATGT<br>CAACGGTCATAAGTTTTCCGTGCGTGGCGAGGGTGAAGGTGACGCAACTAAT<br>GGTAAACTGACGCTGAAGTTCATCTGTACTACTGGTAAACTGCCGGTACCTTG<br>GCCGACTCTGGTAACGACGCTGACTTATGGTGTTCAGTGCTTTGCTCGTTATC<br>CGGACCATATGAAGCAGCATGACTTCTTCAAGTCCGCCATGCCGGAAGGCTA<br>TGTGCAGGAACGCACGATTTCCTTTAAGGATGACGGCACGTACAAAACGCGT<br>GCGGAAGTGAAATTTGAAGGCGATACCCTGGTAAACCGCATTGAGCTGAAAG<br>GCATTGACTTTAAAGAAGATGGCAATATCCTGGGCCATAAGCTGGAATACAAT<br>TTTAACAGCCACAATGTTTACATCACCGCCGATAAACAAAAAAATGGCATTAAA<br>GCGAATTTTAAAATTCGCCACAACGTGGAGGATGGCAGCGTGCAGCTGGCTG<br>ATCACTACCAGCAAAACACTCCAATCGGTGATGGTCCTGTTCTGCTGCCAGA<br>CAATCACTATCTGAGCACGCAAAGCGTTCTGTCTAAAGATCCGAACGAGAAAC<br>GCGATCATATGGTTCTGCTGGAGTTCGTAACCGCAGCGGGCATCACGCATGG<br>TATGGATGAACTGTACAAAATGGCCTGGAAAGGGGAGGTCTTAGCAAACAAT<br>GAGGCCGGACAGGTTACCAGTATCATTTACAATCCAGGGGATGTGATTACGA<br>TTGTTGCGGCTGGTTGGGCGTCTTATGGACCTACTCAGAAGTGGGGACCACA<br>GGGTGATCGTGAACATCCAGATCAGGGGCTGATCTGTCATGACGCATTTTGT<br>GGCGCCCTTGTCATGAAAATTGGCAATAGCGGGACTATTCCAGTAAACACCG<br>GGCTGTTCCGTTGGGTTGCTCCGAACAACGTACAAGGTGCAATTACCTTGAT<br>CTATAATGACGTTCCAGGCACGTATGGAAACAACTCCGGTTCCTTTAGCGTAA<br>ACATTGGCAAGGATCAAAGTTGATAACGACTCAGGCTGCTACTCAAAACTAGC |

| | |
|---|---|
| | ATAACCCCTTGGGGCCTCTAAACGGGTCTTGAGGGGTTTTTTGGGCTTGATAT<br>CGAATTCCTGCAGCCCGGG |
| $P_{T7}$-sfGFP-*lacI* | T7 promoter, ribosomal binding site, sfGFP, *lacI*, T7 terminator |
| | GTCTTCACCTCGAGGATCTTAAGGCTAGAGTAATACGACTCACTATAGGGAGA<br>TGTGGTCTAGACATTCCAGGTTAAGAAGGAGGAAAAAAAAATGCGTAAAGGA<br>GAAGAACTTTTCACTGGTGTCGTCCCTATTCTGGTGGAACTGGATGGTGATGT<br>CAACGGTCATAAGTTTTCCGTGCGTGGCGAGGGTGAAGGTGACGCAACTAAT<br>GGTAAACTGACGCTGAAGTTCATCTGTACTACTGGTAAACTGCCGGTACCTTG<br>GCCGACTCTGGTAACGACGCTGACTTATGGTGTTCAGTGCTTTGCTCGTTATC<br>CGGACCATATGAAGCAGCATGACTTCTTCAAGTCCGCCATGCCGGAAGGCTA<br>TGTGCAGGAACGCACGATTTCCTTTAAGGATGACGGCACGTACAAAACGCGT<br>GCGGAAGTGAAATTTGAAGGCGATACCCTGGTAAACCGCATTGAGCTGAAAG<br>GCATTGACTTTAAAGAAGATGGCAATATCCTGGGCCATAAGCTGGAATACAAT<br>TTTAACAGCCACAATGTTTACATCACCGCCGATAAACAAAAAAATGGCATTAAA<br>GCGAATTTTAAAATTCGCCACAACGTGGAGGATGGCAGCGTGCAGCTGGCTG<br>ATCACTACCAGCAAAACACTCCAATCGGTGATGGTCCTGTTCTGCTGCCAGA<br>CAATCACTATCTGAGCACGCAAAGCGTTCTGTCTAAAGATCCGAACGAGAAAC<br>GCGATCATATGGTTCTGCTGGAGTTCGTAACCGCAGCGGGCATCACGCATGG<br>TATGGATGAACTGTACAAAAAACCAGTAACGTTATACGATGTCGCAGAGTATG<br>CCGGTGTCTCTTATCAGACCGTTTCCCGCGTGGTGAACCAGGCCAGCCACGT<br>TTCTGCGAAAACGCGGGAAAAAGTGGAAGCGGCGATGGCGGAGCTGAATTA<br>CATTCCCAACCGCGTGGCACAACAACTGGCGGGCAAACAGTCGTTGCTGATT<br>GGCGTTGCCACCTCCAGTCTGGCCCTGCACGCGCCGTCGCAAATTGTCGCG<br>GCGATTAAATCTCGCGCCGATCAACTGGGTGCCAGCGTGGTGGTGTCGATG<br>GTAGAACGAAGCGGCGTCGAAGCCTGTAAAGCGGCGGTGCACAATCTTCTC<br>GCGCAACGCGTCAGTGGGCTGATCATTAACTATCCGCTGGATGACCAGGATG<br>CCATTGCTGTGGAAGCTGCCTGCACTAATGTTCCGGCGTTATTTCTTGATGTC<br>TCTGACCAGACACCCATCAACAGTATTATTTTCTCCCATGAAGACGGTACGCG<br>ACTGGGCGTGGAGCATCTGGTCGCATTGGGTCACCAGCAAATCGCGCTGTTA<br>GCGGGCCCATTAAGTTCTGTCTCGGCGCGTCTGCGTCTGGCTGGCTGGCAT<br>AAATATCTCACTCGCAATCAAATTCAGCCGATAGCGGAACGGGAAGGCGACT<br>GGAGTGCCATGTCCGGTTTTCAACAAACCATGCAAATGCTGAATGAGGGCAT<br>CGTTCCCACTGCGATGCTGGTTGCCAACGATCAGATGGCGCTGGGCGCAAT<br>GCGCGCCATTACCGAGTCCGGGCTGCGCGTTGGTGCGGATATCTCGGTAGT<br>GGGATACGACGATACCGAAGACAGCTCATGTTATATCCCGCCGTTAACCACC<br>ATCAAACAGGATTTTCGCCTGCTGGGGCAAACCAGCGTGGACCGCTTGCTGC<br>AACTCTCTCAGGGCCAGGCGGTGAAGGGCAATCAGCTGTTGCCCGTCTCACT<br>GGTGAAAAGAAAAACCACCCTGGCGCCCAATACGCAAACCGCCTCTCCCCGC<br>GCGTTGGCCGATTCATTAATGCAGCTGGCACGACAGGTTTCCCGACTGGAAA<br>GCGGGCAGTAATAACGACTCAGGCTGCTACTCAAAACTAGCATAACCCCTTG<br>GGGCCTCTAAACGGGTCTTGAGGGGTTTTTTGGGCTTGATATCGAATTCCTG<br>CAGCCCGGG |
| $P_{T7}$-sfGFP | T7 promoter, ribosomal binding site, sfGFP, T7 terminator |
| | GTCTTCACCTCGAGGATCTTAAGGCTAGAGTAATACGACTCACTATAGGGAGA<br>TGTGGTCTAGACATTCCAGGTTAAGAAGGAGGAAAAAAAAATGCGTAAAGGA<br>GAAGAACTTTTCACTGGTGTCGTCCCTATTCTGGTGGAACTGGATGGTGATGT<br>CAACGGTCATAAGTTTTCCGTGCGTGGCGAGGGTGAAGGTGACGCAACTAAT<br>GGTAAACTGACGCTGAAGTTCATCTGTACTACTGGTAAACTGCCGGTACCTTG<br>GCCGACTCTGGTAACGACGCTGACTTATGGTGTTCAGTGCTTTGCTCGTTATC<br>CGGACCATATGAAGCAGCATGACTTCTTCAAGTCCGCCATGCCGGAAGGCTA<br>TGTGCAGGAACGCACGATTTCCTTTAAGGATGACGGCACGTACAAAACGCGT<br>GCGGAAGTGAAATTTGAAGGCGATACCCTGGTAAACCGCATTGAGCTGAAAG |

| | |
|---|---|
| | GCATTGACTTTAAAGAAGATGGCAATATCCTGGGCCATAAGCTGGAATACAAT<br>TTTAACAGCCACAATGTTTACATCACCGCCGATAAACAAAAAAATGGCATTAAA<br>GCGAATTTTAAAATTCGCCACAACGTGGAGGATGGCAGCGTGCAGCTGGCTG<br>ATCACTACCAGCAAAACACTCCAATCGGTGATGGTCCTGTTCTGCTGCCAGA<br>CAATCACTATCTGAGCACGCAAAGCGTTCTGTCTAAAGATCCGAACGAGAAAC<br>GCGATCATATGGTTCTGCTGGAGTTCGTAACCGCAGCGGGCATCACGCATGG<br>TATGGATGAACTGTACAAATAATAACGACTCAGGCTGCTACTCAAAACTAGCA<br>TAACCCCTTGGGGCCTCTAAACGGGTCTTGAGGGGTTTTTTGGGCTTGATAT<br>CGAATTCCTGCAGCCCGGG |
| $P_{T7}$-DAGK | T7 promoter, ribosomal binding site, DAGK sequence |
| | GAAATTAATACGACTCACTATAGGGAGACCACAACGGTTTCCCTCTAGAAATA<br>ATTTTGTTTAACTTTAAGAAGGAGATATACCAATGGCCAATAATACCACTGGAT<br>TCACCCGAATTATCAAAGCTGCTGGCTATTCCTGGAAAGGTTTACGCGCTGCA<br>TGGATCAACGAAGCGGCATTCCGTCAGGAAGGCGTAGCGGTATTGTTGGCG<br>GTGGTCATCGCCTGCTGGCTGGATGTGGACGCGATTACCCGCGTGCTGCTTA<br>TCAGCTCCGTGATGCTGGTGATGATTGTGGAAATCCTCAATAGCGCCATCGA<br>AGCAGTGGTTGACCGAATTGGCTCTGAATACCATGAGCTTTCCGGACGCGCA<br>AAAGATATGGGATCCGCTGCGGTGCTGATTGCCATTATCGTCGCCGTGATTA<br>CCTGGTGCATTCTGTTATGGTCGCATTTTGGATAACCGCTGAGCAATAAC |
| $P_{T7}$-sfGFP-DAGK | T7 promoter, ribosomal binding site, sfGFP, DAGK sequence |
| | ATGGCTCGACAGATCTTAAGGCTAGAGTACTTAATACGACTCACTATAGCTAG<br>CATTTAGGTGACACTATAAGCACATCAGCAGGACGCACTGACCGAATTCATTA<br>AAGAGGAGAAAGGTACCATGCATCACCATCACCATCACATGCGTAAAGGAGA<br>AGAACTTTTCACTGGTGTCGTCCCTATTCTGGTGGAACTGGATGGTGATGTCA<br>ACGGTCATAAGTTTTCCGTGCGTGGCGAGGGTGAAGGTGACGCAACTAATGG<br>TAAACTGACGCTGAAGTTCATCTGTACTACTGGTAAACTGCCGGTACCTTGGC<br>CGACTCTGGTAACGACGCTGACTTATGGTGTTCAGTGCTTTGCTCGTTATCCG<br>GACCATATGAAGCAGCATGACTTCTTCAAGTCCGCCATGCCGGAAGGCTATG<br>TGCAGGAACGCACGATTTCCTTTAAGGATGACGGCACGTACAAAACGCGTGC<br>GGAAGTGAAATTTGAAGGCGATACCCTGGTAAACCGCATTGAGCTGAAAGGC<br>ATTGACTTTAAAGAAGATGGCAATATCCTGGGCCATAAGCTGGAATACAATTT<br>TAACAGCCACAATGTTTACATCACCGCCGATAAACAAAAAAATGGCATTAAAG<br>CGAATTTTAAAATTCGCCACAACGTGGAGGATGGCAGCGTGCAGCTGGCTGA<br>TCACTACCAGCAAAACACTCCAATCGGTGATGGTCCTGTTCTGCTGCCAGAC<br>AATCACTATCTGAGCACGCAAAGCGTTCTGTCTAAAGATCCGAACGAGAAAC<br>GCGATCATATGGTTCTGCTGGAGTTCGTAACCGCAGCGGGCATCACGCATGG<br>TATGGATGAACTGTACAAAATGGCCAATAATACCACTGGATTCACCCGAATTA<br>TCAAAGCTGCTGGCTATTCCTGGAAAGGTTTACGCGCTGCATGGATCAACGA<br>AGCGGCATTCCGTCAGGAAGGCGTAGCGGTATTGTTGGCGGTGGTCATCGC<br>CTGCTGGCTGGATGTGGACGCGATTACCCGCGTGCTGCTTATCAGCTCCGTG<br>ATGCTGGTGATGATTGTGGAAATCCTCAATAGCGCCATCGAAGCAGTGGTTG<br>ACCGAATTGGCTCTGAATACCATGAGCTTTCCGGACGCGCAAAAGATATGGG<br>ATCCGCTGCGGTGCTGATTGCCATTATCGTCGCCGTGATTACCTGGTGCATT<br>CTGTTATGGTCGCATTTTGGAGACTACAAAGACGATGACGACAAGTAATAATA<br>ATGGTACCTCTAGAGTCGACCCGGGCGGCCGCAAAAAAAAAAAAAAAAAAA<br>AAAAAAAAAACTAGCATAACCCCTTGGGGCCTCTAAACGGGTCTTGAGGGGT<br>TTTTTGGATCCGGGCTGGCGTAATAGCGAAGAGGCCCGCA |

**Table S3.** Amino acid sequences of proteins purified for this study

| **Protein and size** | **Annotation of features and protein sequence** |
|---|---|
| PIF-sfGFP-ssrA (41.3 kDa) | His$_6$-tag, TEV site, Pif6-tag, sfGFP, ssrA-tag |
| | MHHHHHHENLYFQGGSAGSAGMMFLPTDYCCRLSDQEYMELVFENGQILAKGQ<br>RSNVSLHNQRTKSIMDLYEAEYNEDFMKSIIHGGGGAITNLGDTQVVPQSHVAAA<br>HETNMLESNKHVDRKGEELFTGVVPILVELDGDVNGHKFSVRGEGEGDATNGKL<br>TLKFICTTGKLPVPWPTLVTTLTYGVQCFARYPDHMKQHDFFKSAMPEGYVQER<br>TISFKDDGTYKTRAEVKFEGDTLVNRIELKGIDFKEDGNILGHKLEYNFNSHNVYIT<br>ADKQKNGIKANFKIRHNVEDGSVQLADHYQQNTPIGDGPVLLPDNHYLSTQSVLS<br>KDPNEKRDHMVLLEFVTAAGITHGMDELYKAANDENYALAA |
| PhyB-AviTag (74.1 kDa) | PhyB residues 1-651, AviTag, His$_6$-tag |
| | MVSGVGGSGGGRGGGRGGEEEPSSSHTPNNRRGGEQAQSSGTKSLRPRSNT<br>ESMSKAIQQYTVDARLHAVFEQSGESGKSFDYSQSLKTTTYGSSVPEQQITAYLS<br>RIQRGGYIQPFGCMIAVDESSFRIIGYSENAREMLGIMPQSVPTLEKPEILAMGTD<br>VRSLFTSSSSILLERAFVAREITLLNPVWIHSKNTGKPFYAILHRIDVGVVIDLEPAR<br>TEDPALSIAGAVQSQKLAVRAISQLQALPGGDIKLLCDTVVESVRDLTGYDRVMV<br>YKFHEDEHGEVVAESKRDDLEPYIGLHYPATDIPQASRFLFKQNRVRMIVDCNAT<br>PVLVVQDDRLTQSMCLVGSTLRAPHGCHSQYMANMGSIASLAMAVIINGNEDDG<br>SNVASGRSSMRLWGLVVCHHTSSRCIPFPLRYACEFLMQAFGLQLNMELQLALQ<br>MSEKRVLRTQTLLCDMLLRDSPAGIVTQSPSIMDLVKCDGAAFLYHGKYYPLGVA<br>PSEVQIKDVVEWLLANHADSTGLSTDSLGDAGYPGAAALGDAVCGMAVAYITKR<br>DFLFWFRSHTAKEIKWGGAKHHPEDKDDGQRMHPRSSFQAFLEVVKSRSQPW<br>ETAEMDAIHSLQLILRDSFKESEAAMNSKVVDGVVQPCRDMAGEQGIDELGAGS<br>GSGLNDIFEAQKIEWHEHHHHHH |
| sfGFP-LacI (66.1 kDa) | sfGFP, LacI, His$_6$-tag |
| | MRKGEELFTGVVPILVELDGDVNGHKFSVRGEGEGDATNGKLTLKFICTTGKLPV<br>PWPTLVTTLTYGVQCFARYPDHMKQHDFFKSAMPEGYVQERTISFKDDGTYKTR<br>AEVKFEGDTLVNRIELKGIDFKEDGNILGHKLEYNFNSHNVYITADKQKNGIKANF<br>KIRHNVEDGSVQLADHYQQNTPIGDGPVLLPDNHYLSTQSVLSKDPNEKRDHMV<br>LLEFVTAAGITHGMDELYKKPVTLYDVAEYAGVSYQTVSRVVNQASHVSAKTREK<br>VEAAMAELNYIPNRVAQQLAGKQSLLIGVATSSLALHAPSQIVAAIKSRADQLGAS<br>VVVSMVERSGVEACKAAVHNLLAQRVSGLIINYPLDDQDAIAVEAACTNVPALFLD<br>VSDQTPINSIIFSHEDGTRLGVEHLVALGHQQIALLAGPLSSVSARLRLAGWHKYL<br>TRNQIQPIAEREGDWSAMSGFQQTMQMLNEGIVPTAMLVANDQMALGAMRAITE<br>SGLRVGADISVVGYDDTEDSSCYIPPLTTIKQDFRLLGQTSVDRLLQLSQGQAVK<br>GNQLLPVSLVKRKTTLAPNTQTASPRALADSLMQLARQVSRLESGQHHHHHH |
| sfGFP (27.6 kDa) | sfGFP, His$_6$-tag |
| | MRKGEELFTGVVPILVELDGDVNGHKFSVRGEGEGDATNGKLTLKFICTTGKLPV<br>PWPTLVTTLTYGVQCFARYPDHMKQHDFFKSAMPEGYVQERTISFKDDGTYKTR<br>AEVKFEGDTLVNRIELKGIDFKEDGNILGHKLEYNFNSHNVYITADKQKNGIKANF<br>KIRHNVEDGSVQLADHYQQNTPIGDGPVLLPDNHYLSTQSVLSKDPNEKRDHMV<br>LLEFVTAAGITHGMDELYKHHHHHH |

| | |
|---|---|
| ClpP-msfGFP (50.7 kDa) | ClpP, sfGFP, Mutation to eliminate tendency of GFP to dimerize at high concentrations (11), $His_6$-tag |
| | MSYSGERDNFAPHMALVPMVIEQTSRGERSFDIYSRLLKERVIFLTGQVEDHMANLIVAQMLFLEAENPEKDIYLYINSPGGVITAGMSIYDTMQFIKPDVSTICMGQAASMGAFLLTAGAKGKRFCLPNSRVMIHQPLGGYQGQATDIEIHAREILKVKGRMNELMALHTGQSLEQIERDTERDRFLSAPEAVEYGLVDSILTHRNRKGEELFTGVVPILVELDGDVNGHKFSVRGEGEGDATNGKLTLKFICTTGKLPVPWPTLVTTLTYGVQCFARYPDHMKQHDFFKSAMPEGYVQERTISFKDDGTYKTRAEVKFEGDTLVNRIELKGIDFKEDGNILGHKLEYNFNSHNVYITADKQKNGIKANFKIRHNVEDGSVQLADHYQQNTPIGDGPVLLPDNHYLSTQSKLSKDPNEKRDHMVLLEFVTAAGITHGMDELYKHHHHHH |

**Table S4.** Plasmids created in this study

| Name | Description | Antibiotic | Origin of replication |
|---|---|---|---|
| pTNT-sfGFP-$His_6$ | Expression of sfGFP in *E. coli* BL21(DE3) | Ampicillin | pUC |
| pTNT-sfGFP-LacI-$His_6$ | Expression of sfGFP-LacI in *E. coli* BL21(DE3) | Ampicillin | pUC |
| pTNT-$His_6$-sfGFP-DAGK | Preparation of linear DNA template for TX-TL | Ampicillin | pUC |
| pTNT-ClpP-msfGFP-$His_6$ | Expression of ClpP-msfGFP in *E. coli* BL21(DE3) | Ampicillin | pUC |
| pET11-$His_6$-Pif6-sfGFP-ssrA | Expression of PIF-sfGFP-ssrA in *E. coli* BL21(DE3) | Ampicillin | ColE1 |

### Legends for movies

**Video S1**: Cell-free expression of LecA-sfGFP protein in the presence of droplets. Time-lapse movie of merged brightfield and GFP fluorescence images. LecA-sfGFP partitions into droplets during its synthesis resulting in a gradual increase of droplet fluorescence over time. Movie corresponds to images **Fig. 2*C***.

**Video S2**: Brightfield time-lapse movie of droplet coalescence. Droplets were formed at 0.9 mM GOA and 0.5 mM IGEPAL. Droplets were doped with 25 µM Ni-NTA-DGS and sfGFP-$His_6$ was added to a concentration of 5 µM (fluorescence channel not shown). Images were acquired every 30 s.

**Video S3**: Light-controlled sequestration and release of a PIF-fusion protein by droplets. Time-lapse movie of GFP fluorescence changes of PhyB-loaded droplets, which sequester and release PIF-sfGFP-ssrA in response to red and far red light. The wavelength of the illumination was altered between 660 nm and 740 nm as indicated. This movie is an example of a representative time-lapse movie used to extract fluorescence traces in **Fig 5*B***.
